# Dna2-intrinsic condensation regulates DNA end resection and reveals evolutionary redistribution of condensate grammar

**DOI:** 10.64898/2026.09.16.751900

**Authors:** Chris A. Jordan, Jonas Lehar, Brandon J. Payliss, Guillaume Laflamme, Zhongle Liu, Juliana The, Janet N. Y. Chan, Tamari Chkuaseli, Razq Hakem, Leah E. Cowen, Haley D. M. Wyatt, Karim Mekhail

## Abstract

Eukaryotic cells commonly use biomolecular condensation of DNA double-strand break (DSB) repair scaffolds and signaling assemblies to organize repair reactions in space and time. Yet whether DSB end-processing enzymes themselves encode tunable phase separation that potentiates resection remains unclear. Here we show that the long-range resection enzyme Dna2 forms liquid-like condensates through an intrinsically disordered region that is necessary and sufficient for phase separation and catalytic enhancement in *Saccharomyces cerevisiae*. Dna2 condensates concentrate DNA substrates and enhance end processing, whereas disrupting condensate formation impairs repair kinetics, checkpoint signaling, and chromosome stability. Grafting of the heterologous intrinsically disordered region of human FUS partly rescues condensate formation and function, and Cdk1-dependent phosphorylation sites tune condensate stability and enzymatic output in cis and in trans. Machine-learning-based analysis reveals that condensation-promoting features of fungal Dna2 are shared with a restricted set of human DNA2-associated resection regulators. Together, these findings define phosphorylation-tuned, enzyme-intrinsic phase separation as an organizational principle of DSB end resection while supporting a model in which condensation-promoting features are redistributed among factors operating within conserved genome maintenance pathways.

## Introduction

DNA double-strand breaks (DSBs) threaten genome stability and cell survival. The repair of DSBs relies on evolutionarily conserved pathways, including homologous recombination (HR) and non-homologous end joining (NHEJ), coordinated by a broad DNA damage response (DDR)^1,2^. A key determinant of repair pathway choice is DNA end resection, which generates long 3′ single-stranded DNA (ssDNA) intermediates that commit repair to HR^2^. By licensing HR and amplifying checkpoint signaling, end resection acts as a major regulatory node in genome maintenance while requiring tight control of highly potent DNA-processing enzymes.

In budding yeast, HR initiation involves checkpoint activation followed by a two-stage resection program. Short-range processing is mediated by the Mre11-Rad50-Xrs2 complex together with Sae2^3–7^, providing an entry point for long-range resection by either the exonuclease Exo1 or the nuclease-translocase Dna2 acting with the RecQ helicase Sgs1^8–14^. While Exo1 preferentially processes accessible DNA ends, Dna2 can resect more structurally complex or protein-bound DNA substrates^5,9^. In both yeast and mammals, long-range resection generates RPA-coated ssDNA that promotes checkpoint activation and Rad51-mediated strand invasion^2^.

Beyond enzymatic chemistry, increasing evidence indicates that DNA repair pathways are spatially organized through biomolecular condensation. Several DNA repair scaffolds, signaling factors, and chromatin-associated proteins form condensates that concentrate repair factors and organize repair reactions in space and time^1,15–39^. Intrinsically disordered regions (IDRs) frequently contribute to such assemblies through multivalent and dynamic interactions. However, whether DSB end-processing enzymes themselves encode tunable condensation that directly potentiates catalytic function remains unknown. More broadly, how condensation-promoting sequence grammar is organized and evolutionarily distributed across conserved molecular pathways remains poorly understood.

Here, using complementary cellular, biochemical, and computational approaches, we show that the long-range resection enzyme Dna2 intrinsically forms liquid-like condensates through an N-terminal intrinsically disordered region that directly potentiates DNA end processing. We further identify phosphorylation as a regulator of Dna2 condensate properties and uncover patterns consistent with evolutionary redistribution of condensation-promoting sequence features across conserved DNA repair pathways.

## Results

### Intrinsic disorder identifies liquid-like Dna2

To gain insights into the broader potential of DNA damage response proteins (DDRome) to intrinsically encode condensation, we predicted protein disorder across the *S. cerevisiae* proteome using two independent computational approaches^40–42^ and compared disorder scores between DDR proteins and the remainder of the proteome. DDR proteins were, on average, significantly more disordered, both in total predicted disorder length and the fraction of the protein occupied by predicted disorder (Fig. 1a,b; Supplementary Fig. 1a-c). The HR scaffold Rad52 and the ssDNA-binding factor Rfa1 displayed high and low disorder scores, respectively (Supplementary Fig. 1d), consistent with the known ability of Rad52, but not Rfa1, to phase separate in yeast^21,24,43^.

**Fig. 1.**
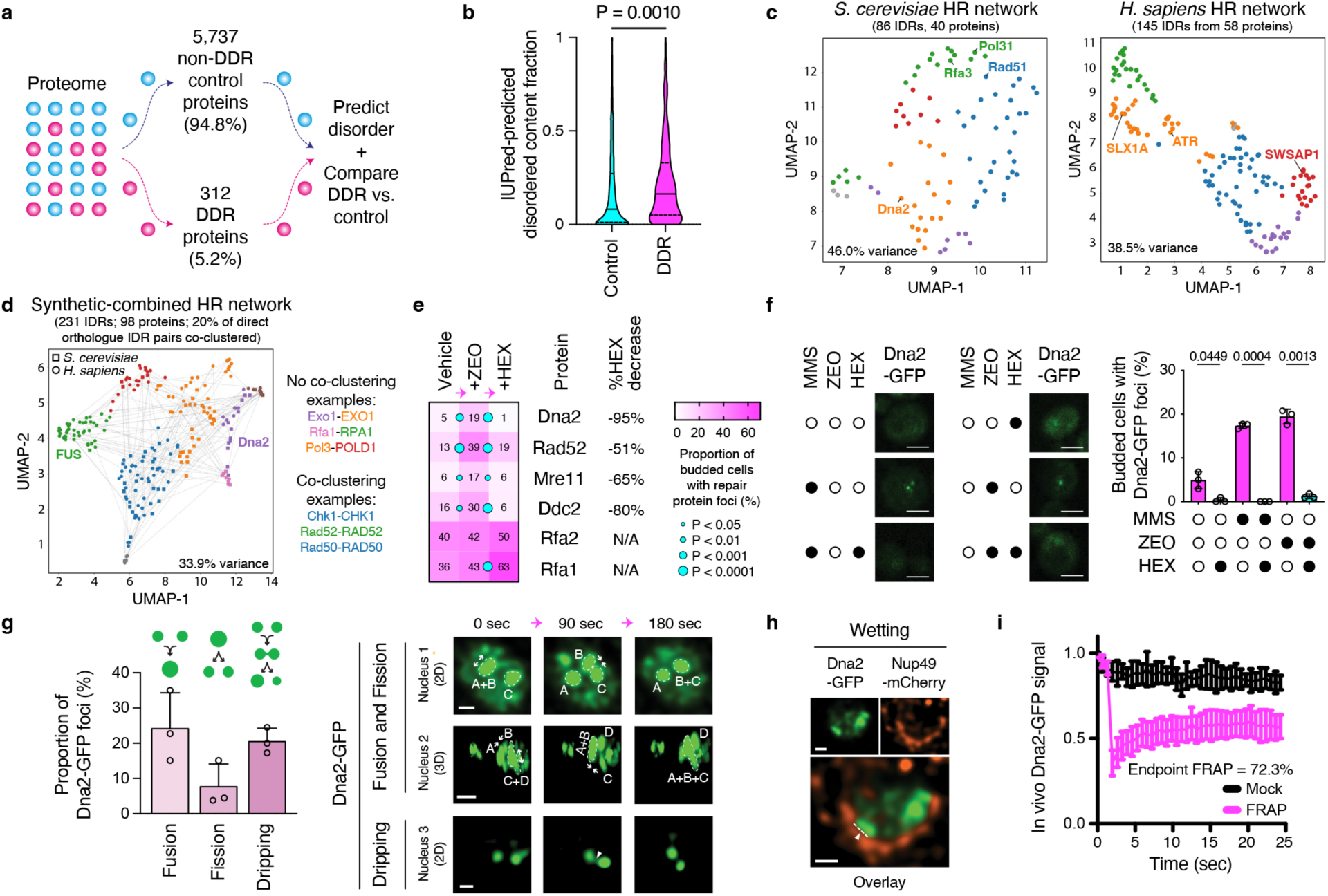
Intrinsic disorder identifies Dna2 as a liquid-like repair factor. **a**, Schematic of in silico intrinsic disorder analysis across the *S. cerevisiae* proteome. **b**, Disordered fraction of DNA damage response (DDR) proteins and control proteins predicted by IUPred. Data are shown as the mean ± interquartile range; *n* = 5,737 control proteins and 312 DDR proteins; two-tailed unpaired t-test. **c**-**d**, UMAP projections from machine-learning comparative analyses of condensate grammar of intrinsically disordered regions (IDRs) within homologous recombination (HR) proteins. Points represent individual IDRs labeled by hierarchical clustering assignment. Projections are separated by species (*S. cerevisiae*, left; *H. sapiens*, right) in (**c**), with lines connecting direct yeast-human orthologues in (**d**). Conserved pathway components with disorder in only one of the two species are indicated (**c**), and IDRs from proteins particularly relevant to this study are shown on the plot (**d**). **e**, Cells expressing GFP-tagged DDR proteins were treated with zeocin (ZEO, 50 μg.ml^-1^, 1 hr) to induce repair foci before testing their sensitivity to 1,6-hexanediol (HEX, 5%). Heatmap shows the mean; *n* = 3 independent biological replicates, two-way ANOVA with Dunnett’s test. HEX-dependent percent decreases are also indicated. **f**, Dna2-GFP foci induced by a one-hour treatment with methyl methanesulfonate (MMS, 0.03%) or ZEO (ZEO, 50 μg.ml^-1^) and their sensitivity to HEX. Representative images (left) and mean quantification (right) are shown. Data show the mean ± s.d.; *n* = 3 independent biological replicates, two-tailed unpaired *t*-test with Welch’s correction; scale bars, 5 μm. **g**, Quantification (left) and representative images (right) of fusion, fission, and dripping of MMS-induced Dna2-GFP foci within the imaging time window. Data show the mean + s.d.; *n* = 3 independent biological replicates; scale bars, 0.5 μm for two-dimensional (2D) and 2 μm for three-dimensional (3D) images. **h**, Surface wetting of MMS-induced Dna2-GFP foci at the Nup49-mCherry-marked nuclear envelope. Scale bars, 0.5 μm. **i**, Fluorescence recovery after photobleaching (FRAP) analysis of MMS-induced Dna2-GFP foci. Data show the mean ± s.d.; *n* = 29 cells from 3 independent biological replicates.

We next asked whether predicted intrinsically disordered regions (IDRs) cluster according to their sequence features (collectively known as condensate grammar) within HR or NHEJ networks in *S. cerevisiae* and *H. sapiens*. To address this question, we performed comparative machine-learning-based analyses^44^ considering 90 condensate-associated organizational features from predicted disordered regions across HR and NHEJ proteins. The resulting UMAP projections revealed that conserved repair pathways exhibit species-specific architectures of condensate grammar networks (Fig. 1c; Supplementary Fig. 1e). Most notably, the central resection factor Dna2, as well as Rad51 and Rfa3, contained predicted disordered regions in *S. cerevisiae* while their human orthologues did not (Fig. 1c). In addition, using yeast-human comparative analyses, amongst IDR pairs connecting yeast and human orthologous proteins, only 20% and 22% co-clustered when assessing HR and NHEJ, respectively, suggesting that these pairs frequently exhibit divergent condensate grammar (Fig. 1d; Supplementary Fig. 1f). Together, these results are consistent with extensive evolutionary reorganization of condensate-promoting sequence features across conserved DNA repair pathways.

To experimentally test the ability of predicted disorder to promote protein assemblies in cells exposed to DNA damage, we endogenously GFP-tagged several *S. cerevisiae* DDR proteins spanning a range of predicted disorder levels (Supplementary Fig. 1d,g). All tagged strains retained resistance to the DNA-damaging agents zeocin and methyl methanesulfonate (MMS), indicating preserved protein function (Supplementary Fig. 1h). We next examined the sensitivity of DNA damage-induced repair foci to the aliphatic alcohol 1,6-hexanediol, which disrupts many phase-separated assemblies^45,46^. Amongst the proteins tested, Dna2-GFP repair foci displayed pronounced sensitivity to 1,6-hexanediol following zeocin or MMS treatment (Fig. 1e,f; Supplementary Fig. 1i; unless otherwise indicated, cells were treated with 0.03% (w/v) MMS or 50 μg.ml^-1^ zeocin for 1 h). In addition, the presence of a predicted disordered and structurally uncharacterized N-terminal region within *S. cerevisiae* Dna2 (Supplementary Fig. 2a,b), together with previous observations associating this region with chromosome maintenance^47–49^, identified Dna2 as a compelling candidate for mechanistic analysis of enzyme-intrinsic condensation. Moreover, because Dna2 functions as a central long-range resection enzyme in both yeast and human cells, divergence in its predicted disorder architecture across these species (Fig. 1c) offered an opportunity to investigate how condensation-promoting sequence features may evolve within the conserved HR pathway.

We next asked whether Dna2 foci display dynamic properties characteristic of liquid-like biomolecular condensates in vivo. Condensate-like foci typically display several key behaviors over time in live cells^21,22,50^. These behaviors include fusion, fission, and dripping, as observed for Dna2-GFP foci using single-live-cell time-lapse imaging (Fig. 1g; Supplementary Fig. 2c,d). Droplets quickly relaxed to a spherical shape following fusion (Supplementary Fig. 2c). In addition, Dna2-GFP foci contacting the Nup49-mCherry-marked nuclear envelope were flattened and conformed to the envelope’s shape (Fig. 1h), consistent with the surface wetting behavior often exhibited by phase-separated compartments^21,22^. Moreover, Dna2-GFP showed rapid signal recovery in fluorescence recovery after photobleaching (FRAP) experiments (Fig. 1i; Supplementary Fig. 2e; Supplementary Video 1), concordantly with liquid-like behavior^21,22,51^. By comparison, the microtubule-marking GFP-Tub1 showed smaller and slower recovery in FRAP experiments (Supplementary Fig. 2f), serving as a negative control as expected^21^. Treating cells with sorbitol alone, which promotes phase separation^52^ without inducing DNA damage, increased Dna2-GFP foci formation (Supplementary Fig. 2g-i). Furthermore, Dna2-GFP foci were markedly less sensitive to 2,5-hexanediol, a structural isomer used as a negative or reduced activity control for aliphatic alcohol-mediated condensate disruption^45,53^, than to 1,6-hexanediol (Supplementary Fig. 2j). Together, our observations indicate that intrinsic disorder is overrepresented among DDR proteins and that the long-range resection enzyme Dna2 exhibits multiple behaviors consistent with phase-separated, liquid-like biomolecular condensates.

### Dna2 IDR forms condensates and enhances catalysis in vitro

Next, we assessed the phase-separation capacity of the eGFP-tagged N-terminal IDR of *S. cerevisiae* Dna2 (IDR^Dna2^). In vitro, *S. cerevisiae* IDR^Dna2^ purified from *Escherichia coli* phase-separated from the buffer and formed detectable droplets in the presence of the crowding agent PEG, which facilitates condensate formation (Fig. 2a,b; Supplementary Fig. 3a). Droplet size correlated with protein concentration (Fig. 2c,d). Phase separation was readily detected across most mid-range salt concentrations tested, indicating an optimal range from over 25 mM to below 200 mM (Fig. 2e). IDR^Dna2^ droplets commonly fused with one another (Fig. 2f; Supplementary Video 2) and attracted Cy3-labeled ssDNA from the surrounding solution, concentrating the labeled nucleic acid within the droplets (Fig. 2g). The droplets were sensitive to 1,6-hexanediol (Fig. 2h) and exhibited rapid and near-complete fluorescence recovery in FRAP assays (Fig. 2i,j; Supplementary Video 3). Together, these results indicate that the N-terminal region of Dna2 exhibits multiple behaviors consistent with liquid-like biomolecular condensates.

**Fig. 2.**
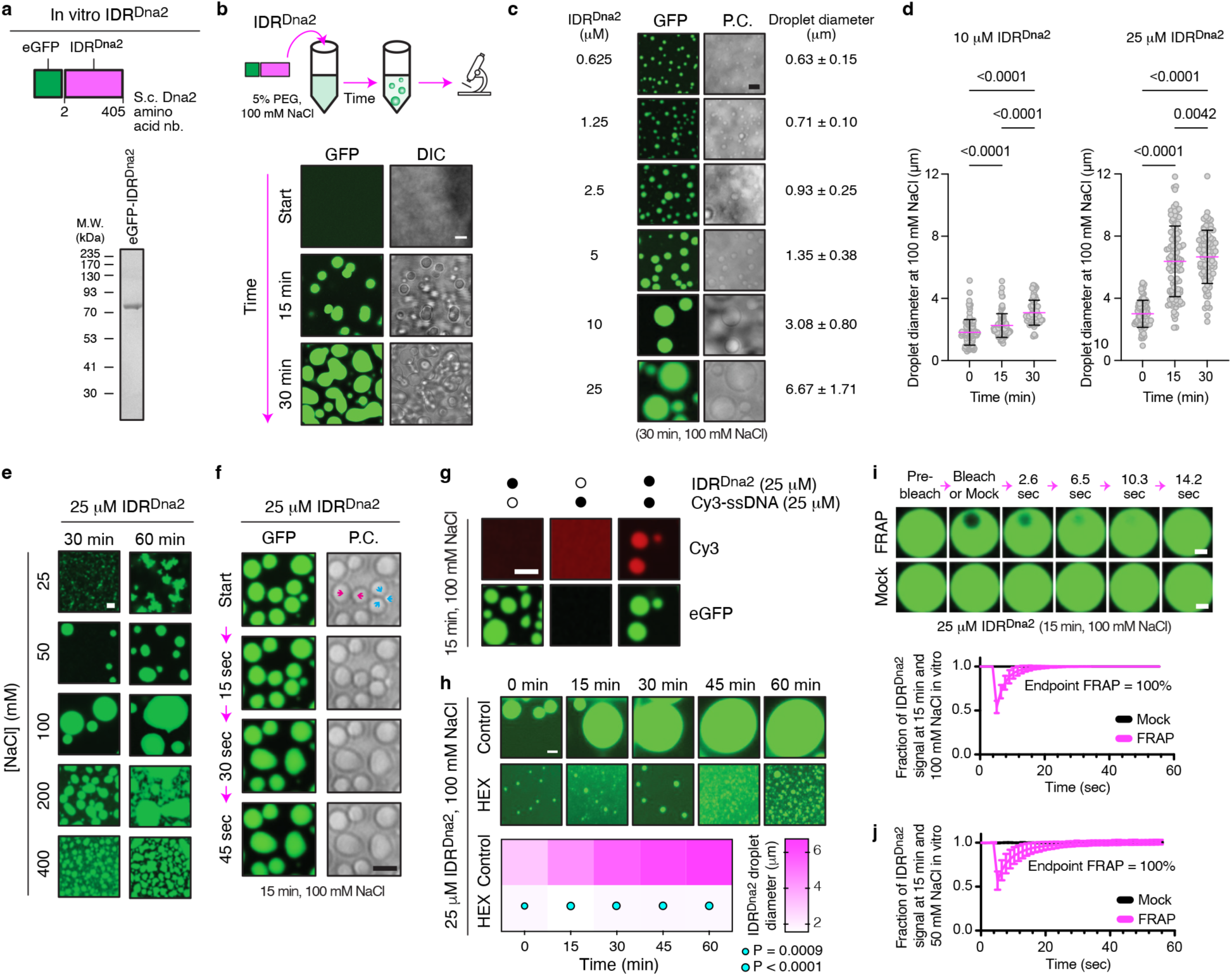
An intrinsically disordered region of Dna2 forms DNA-recruiting condensates. **a**, IDR^Dna2^ schematic (top) and recombinant protein purified from *E. coli* (bottom). **b**, Experimental setup (top) and representative images of IDR^Dna2^ (bottom) in vitro. Scale bar, 5 μm. **c**, Dependence of IDR^Dna2^ droplet size on protein concentration. P.C., phase contrast; scale bar, 5 μm. **d**, Time- and concentration-dependence of IDR^Dna2^ droplet size. Data show the mean ± s.d.; from left to right, *n* = 111, 87, 62, 80, 106, and 89 droplets from 3 independent cell-free replicates, two-way ANOVA with Tukey’s test. **e**, Salt dependence of IDR^Dna2^ droplet formation. Scale bar, 2 μm. **f**, Fusion of IDR^Dna2^ droplets (arrowheads). Scale bar, 2 μm. **g**, IDR^Dna2^ droplets concentrate ssDNA from solution. Scale bar, 2 μm. **h**, Representative images (top) and heatmap (bottom) showing sensitivity of IDR^Dna2^ droplets to 1,6-hexanediol (HEX). Heatmap shows the mean; from left to right, *n* = 30, 40, 42, 34, and 35 droplets for control and *n* = 30, 24, 31, 37, and 42 for HEX, from 3 independent cell-fee replicates, two-way ANOVA with Sidak’s test. Scale bar, 2 μm. **i**,**j**, Representative images (**i**, top) and quantification (**i**, bottom; **j**) of fluorescence recovery after photobleaching (FRAP) of IDR^Dna2^ droplets at 100 mM salt (**i**) and 50 mM salt (**j**). Quantification shows the mean ± s.d.; *n* = 20 (**i**, FRAP), 12 (**i**, Mock), 10 (**j**, FRAP), and 10 (**j**, Mock) droplets from 3 independent cell-free replicates. Scale bar, 2 μm.

Since IDR^Dna2^ droplets have the intrinsic ability to recruit ssDNA (Fig. 2g), we next asked whether these assemblies can reorganize DNA in the presence of binding partners. In the presence of purified yeast RPA complex (yRPA)^21^, the IDR^Dna2^ formed enlarged condensates (Supplementary Fig. 3b,c), and upon subsequent ssDNA addition, these droplets underwent a time-dependent transition in which fluorescent ssDNA reorganized into transient filament-like structures that extended from or merged into the condensates (Supplementary Fig. 3d-f; Supplementary Video 4). Thus, IDR^Dna2^ condensates can promote higher-order organization of ssDNA under defined in vitro conditions.

We next assessed whether intrinsic phase separation affects Dna2’s ability to process a 5′-ssDNA flap substrate in vitro. We first compared the nuclease activity of full-length Dna2 or the IDR-less dna2Δ405 purified from *S. cerevisiae* by measuring the proportions of all reaction products (Pt) and more advanced reaction products (Pa) (Fig. 3a-c). Such differential product-based measurement approaches assess reaction kinetics more comprehensively^54^. Compared to Dna2, dna2Δ405 exhibited significantly lower activity as measured by the Pt or Pa fraction (Fig. 3d-f; compare columns 1 and 5 within each heatmap). In addition, Pt-measured activity increased for both Dna2 and dna2Δ405 in the presence of the crowding agent PEG (Fig. 3d,e; compare heatmap columns 1 to 3 and 5 to 7), and longer reactions confirmed that non-PEG-treated samples require more time but can eventually reach activity levels similar to those of their PEG-treated counterparts (Supplementary Fig. 4a-c). Although PEG enhanced processing by both proteins, dna2Δ405 remained measurably less active than full-length Dna2 by the experimental endpoint even under crowding conditions (Fig. 3d-f; Supplementary Fig. 4a-c), consistent with the IDR conferring catalytic potentiation beyond non-specific macromolecular crowding. Notably, 1,6-hexanediol disproportionately suppressed formation of advanced products by full-length Dna2 relative to dna2Δ405 regardless of PEG (Fig. 3d,f), indicating that sustained catalytic progression depends on an IDR-dependent higher-order state that is selectively sensitive to aliphatic alcohol-based disruption. Together, these observations indicate that Dna2 harbors an N-terminal IDR that is sufficient for phase separation and enhances catalysis in vitro. Of note, these reactions were conducted without the addition of other DNA repair factors, such as RPA, Sgs1, or the Top3–Rmi1 complex^2^, consistent with Dna2 possessing an intrinsic capacity to promote higher-order organization that enhances end processing. While PEG facilitates assembly under these in vitro conditions, concordance between in vitro (Fig. 2 and 3) and cellular (Fig. 1) observations suggests the physiological relevance of Dna2-dependent higher-order organization.

**Fig. 3.**
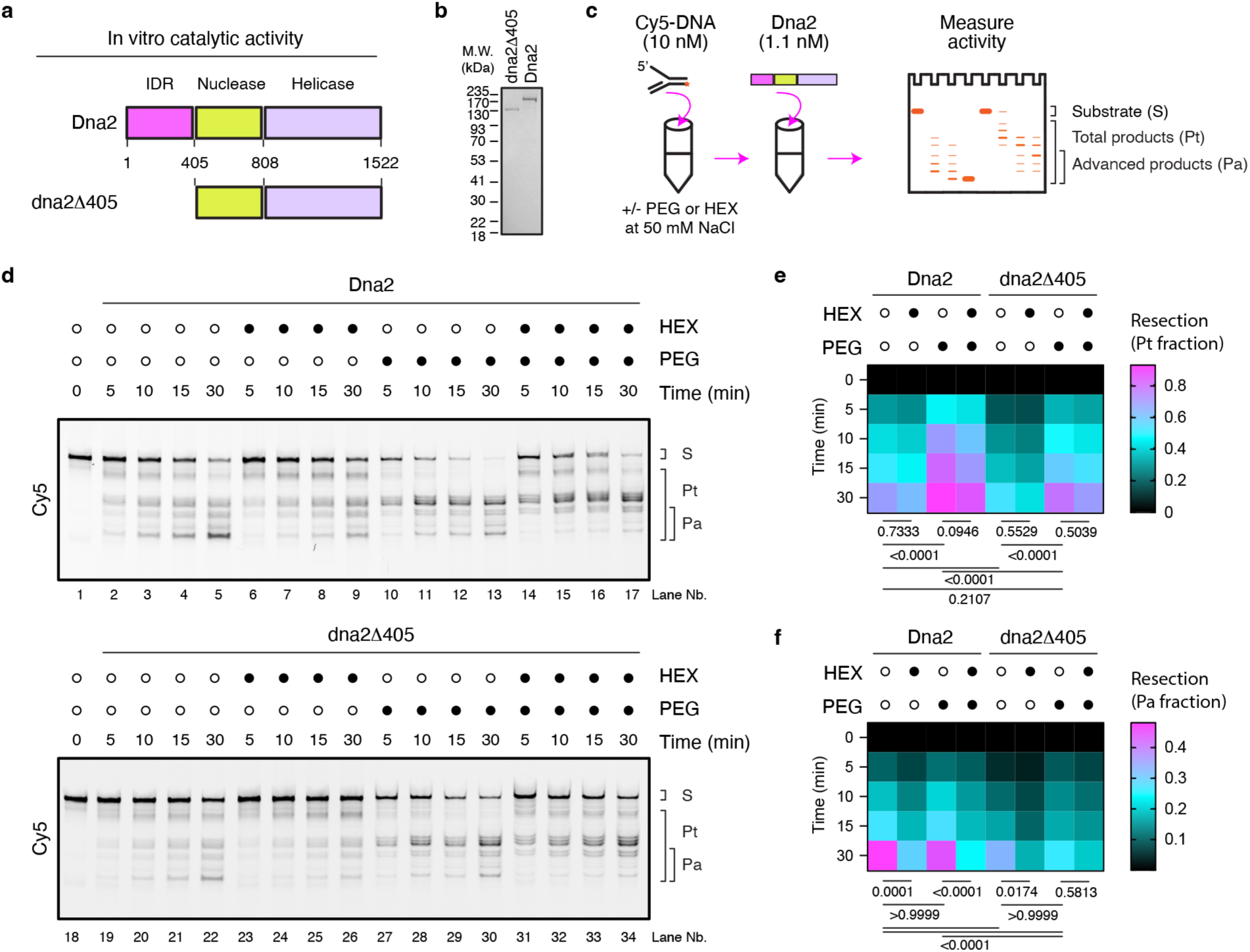
Condensation potentiates Dna2 function in vitro. **a**,**b**, Schematics (**a**) and representative gel (**b**) of full-length Dna2 and IDR-less Dna2 (dna2Δ405) purified from *S. cerevisiae*. **c**, Experimental design of the in vitro nuclease activity assay used to compare full-length and IDR-less Dna2. The Cy5-labelled 5’-ssDNA flap substrate (S), total products (Pt), and advanced products (Pa) are illustrated. **d**-**f**, Representative gels (**d**) and quantification of full-length or IDR-less Dna2 activity based on the Pt fraction (**e**) or Pa fraction (**f**) in the presence or absence of 5% polyethylene glycol (PEG), 5% 1,6-hexanediol (HEX), or both. Data show the mean from *n* = 3 independent cell-free replicates; two-way ANOVA with Tukey’s test (**e**,**f**).

### Genome stability by natural or synthetic Dna2 condensates

We then tested whether removing IDR^Dna2^ reduces Dna2-eGFP’s ability to form foci during DSB repair in live cells. However, the IDR^Dna2^ harbors a nuclear localization signal. Therefore, we replaced the IDR^Dna2^ with a nuclear localization signal alone (Fig. 4a), confirming that both the full-length and IDR-less proteins localize to the nucleus (Supplementary Fig. 5a,b). Importantly, the IDR was required for the protein’s ability to form foci (Fig. 4b) and for the ability of cells to survive exposure to MMS and especially zeocin (Supplementary Fig. 5c). To assess the impact of losing Dna2 or its IDR on overall chromosome stability, we separated whole chromosomes using contour-clamped homogeneous electric field (CHEF) electrophoresis. This analysis revealed that Dna2-deficient cells that were untreated, or especially subjected to exogenous damage, exhibit increased chromosomal smearing and a decrease in chromosome band intensity following recovery (Supplementary Fig. 5d; lanes 1-6). Note that all *dna2Δ* cells were of a *pif1-m2* background to ensure cell viability^9,11,48,55^. Complementation with full-length *DNA2* partly restored wild-type patterns (Supplementary Fig. 5d; lanes 1-9), whereas expression of dna2Δ405 resulted in chromosomal instability both before and after exposure to exogenous damage (Supplementary Fig. 5d; lanes 10-12). Relative to *DNA2*-proficient cells, cells expressing only dna2Δ405 (Fig. 4c) or lacking any Dna2 protein (Supplementary Fig. 5e) displayed delayed DSB repair kinetics as measured by phosphorylated H2A (γH2A), which is analogous to the human γH2AX histone variant^56^. Dna2-mediated DSB end resection amplifies the DNA damage checkpoint, whose activation can be assessed by the proportion of phosphorylated and higher-migrating bands of Rad53, the CHK2 tumor suppressor in mammals^57^. As expected, *dna2Δ* cells displayed a lower degree of Rad53 activation following exposure to zeocin (Supplementary Fig. 5f; lanes 1-4). Similarly, *dna2Δ* cells complemented with *dna2Δ405* displayed a lower degree of Rad53 activation compared to when the cells were complemented with wild-type *DNA2* (Supplementary Fig. 5f; lanes 5-8). Together, these data indicate that IDR^Dna2^ is required for Dna2 focus formation and supports checkpoint amplification, genome repair, and cell survival.

**Fig. 4.**
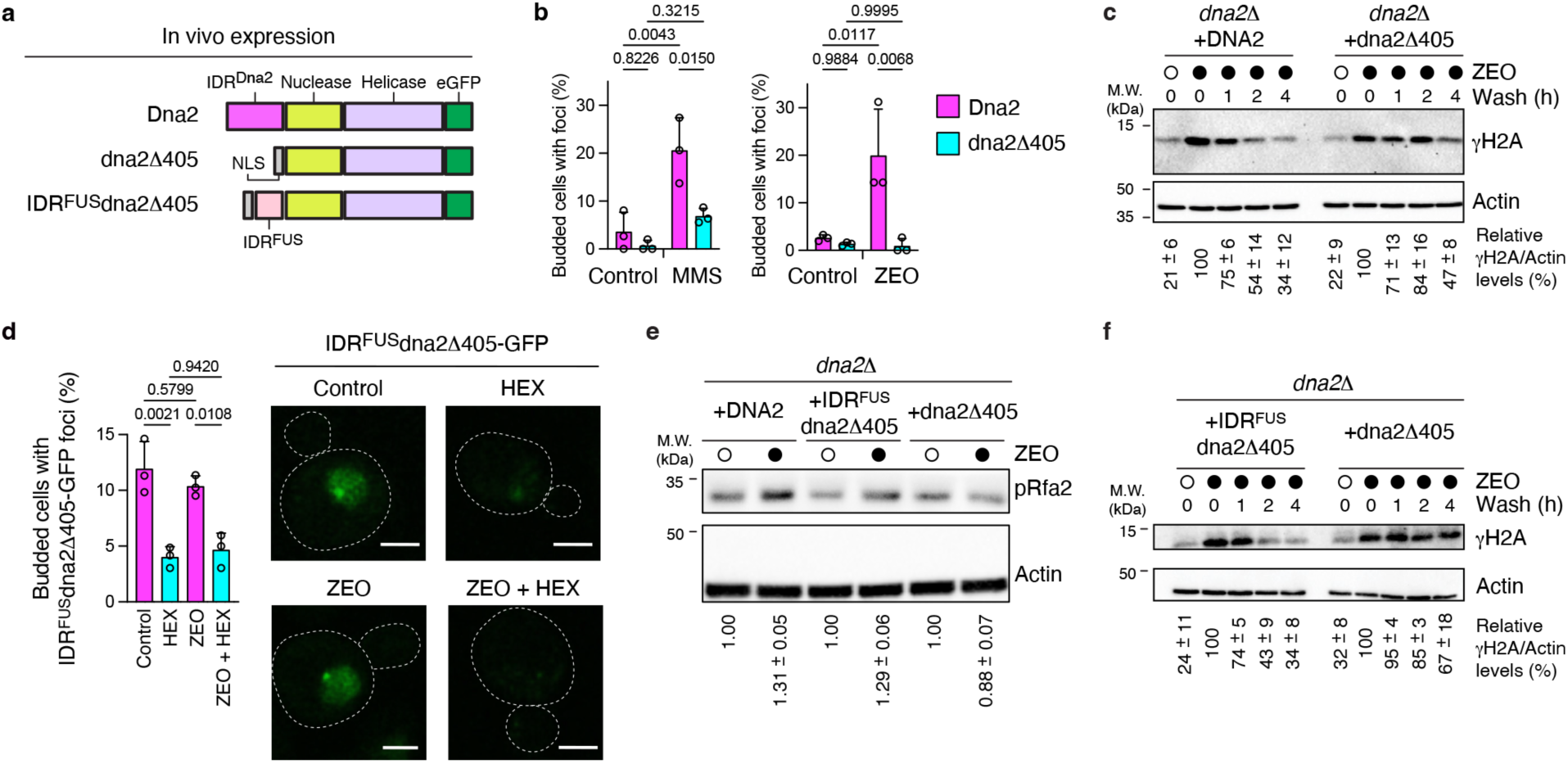
EVicient resection and genome stability through cellular Dna2 condensates. **a**, Schematics of full-length Dna2 and IDR-less Dna2 without or with fusion to IDR^FUS^. NLS, nuclear localization signal. **b**,**c**, Effect of IDR^Dna2^ removal on methyl methanesulfonate (MMS)- or zeocin (ZEO)-induced repair foci formation (**b**) and γH2A-marked genome instability (**c**). Data (**b**) show the mean + s.d. from *n* = 3 independent biological replicates, each with over 30 cells, two-way ANOVA with Tukey’s test. Data (**c**) below immunoblots show the mean ± s.e.m. from *n* = 4 independent biological replicates. **d**, Quantification (left) and representative images (right) showing foci formation by IDR-less Dna2 fused to IDR^FUS^ in the presence of ZEO, 1,6-hexanediol (HEX), or both. Data show the mean + s.d.; *n* = 3 independent biological replicates, each with over 30 cells, two-way ANOVA with Tukey’s test. Scale bar, 2 μm. **e**,**f**, Immunoblots showing the effect of fusing IDR^FUS^ to IDR-less Dna2 on ZEO-induced phosphorylated Rfa2 (pRfa2) levels (**e**) and γH2A-marked genome instability (**f**). Data below immunoblots show the mean ± s.e.m. from *n* = 5 (**e**) or *n* = 3 (**f**) independent biological replicates.

To more directly test this model, we asked whether fusing dna2Δ405 to the heterologous IDR domain of the human protein FUS (IDR^FUS^; Fig. 1d)^17,58^ would rescue the ability of dna2Δ405 to form foci and promote DNA repair (Fig. 4a; Supplementary Fig. 5a). In live untreated cells or zeocin-treated cells, the eGFP-tagged IDR^FUS^-dna2Δ405 formed nuclear foci sensitive to 1,6-hexanediol (Fig. 4d). Note that unlike wild-type Dna2, these assemblies were readily detectable prior to exogenous damage induction. In addition, following exposure to zeocin, *dna2Δ* cells (Supplementary Fig. 5g) or IDR^Dna2^-less cells (Fig. 4e; compare lanes 5-6 to 1-2) showed decreased accumulation of phosphorylated Rfa2 (pRfa2), which provides a measure of functionally competent resection generating ssDNA-RPA sufficient to engage Mec1/Tel1 signaling^21,59^. Moreover, fusing IDR^FUS^ to dna2Δ405 rescued zeocin-dependent increases in pRfa2 levels (Fig. 4e), restoring the ability of cells to activate resection-dependent checkpoint signaling (Supplementary Fig. 5h) and more rapidly resolve DSB levels (Fig. 4f). These observations are consistent with a model in which heterologous IDR^FUS^ promotes constitutive higher-order assembly that subsequently engages DNA lesions following recruitment through Sgs1. Indeed, the proportion of cells with Rfa1-marked DNA damage sites colocalizing with IDR^FUS^-dna2Δ405 robustly decreased upon knocking out Sgs1 (Supplementary Fig. 5i), in agreement with Sgs1’s ability to recruit Dna2 to DSBs^5,9^. We note that although IDR^FUS^ partially restored early repair-associated functions, it did not recapitulate the regulated damage-induced dynamics of native Dna2 assemblies (Fig. 4d) or fully rescue long-term genome stability at the concentration tested (Supplementary Fig. 5j). This agrees with our comparative analyses indicating that the condensate grammar of IDR^FUS^ differs from that of IDR^Dna2^. This further suggests that condensate-forming capacity alone is important but insufficient for complete functional restoration and that sequence-encoded organizational features contribute to native Dna2 function. Our results are also consistent with a division of labor where Sgs1 promotes Dna2 engagement at lesions while Dna2 condensation sustains long-range processing.

### Condensate tuning potentiates resection

The ability of *S. cerevisiae* Dna2 to phase-separate upon DSB induction suggests that upstream DDR signaling controls this condensation. One candidate regulatory mechanism may involve the Cdk1-dependent phosphorylation of key Dna2 N-terminal residues^48,60^. To test this hypothesis, we generated and purified eGFP-tagged *S. cerevisiae* IDR^Dna2^ variants harboring mutations at Cdk1-dependent phosphorylation sites (Fig. 5a (top); Supplementary Fig. 6a). This yielded the wild-type IDR^Dna2^, its phosphomimetic IDR-3D^Dna2^ (carrying T4D, S17D, and S237D mutations), and phosphonull IDR-3A^Dna2^ (carrying T4A, S17A, and S237A mutations).

**Fig. 5.**
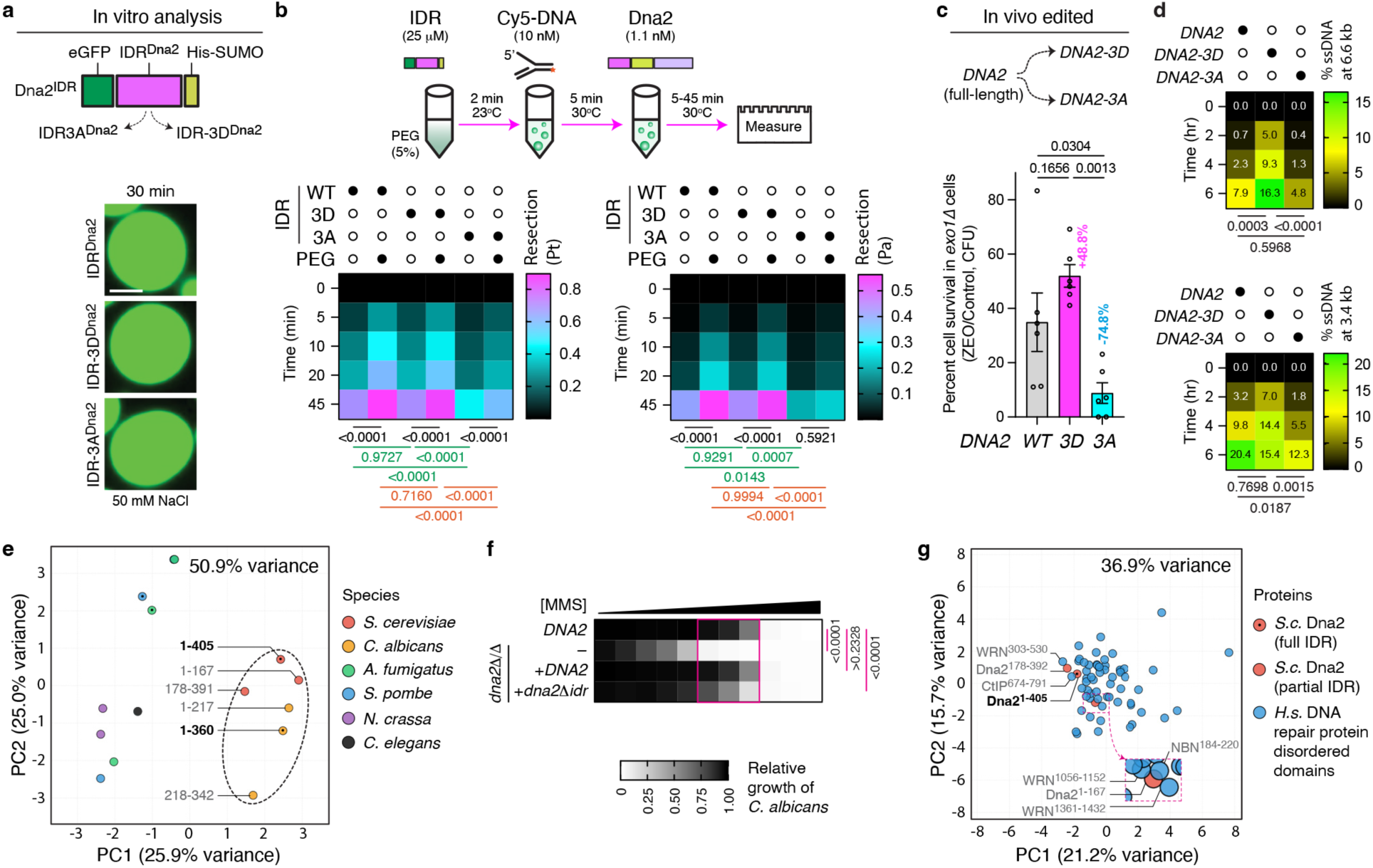
Dna2 condensates are tunable and reveal evolutionary redistribution of condensate grammar. **a**, Schematic (top) and representative droplet images (bottom) of the IDR of Dna2 (IDR^Dna2^) and its phosphomimetic (IDR-3D^Dna2^) and phosphonull (IDR-3A^Dna2^) variants. **b**, Experimental design (top) and total (Pt) or advanced (Pa) activity (bottom) testing the in-trans effect of preformed IDR^Dna2^ droplet variants on the processing of a Cy5-labeled 5’-ssDNA flap by full-length Dna2. Data show the mean; *n* = 3 independent cell-free replicates; two-way *ANOVA* with Tukey’s test. **c**, Schematic of full-length Dna2 and its phosphomimetic (Dna2-3D) and phosphonull (Dna2-3A) variants (top) and their effect on cell survival in the presence of 10 μg.ml^-1^ ZEO for 5 days (bottom). Data show the mean ± s.e.m., *n* = 3 independent biological replicates (each with 2 technical replicates), two-way ANOVA with Tukey’s test. **d**, Impact of Dna2, Dna2-3D, and Dna2-3A on resection-indicating ssDNA 3.4 or 6.6 kb from a single DSB 0 to 6 hrs after its induction in *exo1Δ* background cells. Data show the mean, *n* = 4 (*DNA2-3D* and *DNA2-3A*) or *n* = 3 (*DNA2*) independent biological replicates, each from three technical replicates, two-way ANOVA with Tukey’s test. **e**, Principle component analysis (PCA) projections from machine-learning-based comparative condensate grammar analysis of the IDR in Dna2 from *S. cerevisiae* and other fungi, with *C. elegans* control. **f**, Effect of removing the IDR from *Candida albicans* Dna2 on sensitivity to methyl methanesulfonate (MMS). Both *DNA2* alleles were knocked out (*dna2Δ/Δ*) before complementation with each full-length or IDR-less *DNA2* allele. Data show the mean, *n* = 6 (*dna2Δ/Δ* with each of 2 full-length or 2 IDR-less alleles, 3 clones each) or *n* = 2 independent biological cultures (DNA2 and *dna2Δ/Δ*), two-way *ANOVA* with Dunnett’s test. **g**, PCA projections from machine-learning-based assessment of the condensate grammar similarity of *S. cerevisiae* IDR^Dna2^ and IDRs predicted within 25 major human DNA repair proteins (details in Supplementary Fig. 10c).

We next tested whether the different IDRs create an environment conducive to Dna2-dependent function in vitro. To avoid confounding effects of modifying IDR^Dna2^ within the full length Dna2 on its catalytic domains, we first preassembled IDR droplets and then added unmodified full-length Dna2 to measure nuclease activity (Fig. 5b (top)). Under 50 mM NaCl, despite similar droplet average diameter and 5’-ssDNA flap substrate recruitment, IDR^Dna2^ and IDR-3D^Dna2^ droplets were more spherical and supported end processing more effectively as compared to IDR-3A^Dna2^ droplets in vitro (Fig. 5a,b; Supplementary Fig. 6b-e). These findings indicate that condensate material properties, rather than size or substrate partitioning alone, govern catalytic potentiation. Of particular note, under these conditions, the addition of Dna2 in the presence of PEG induced Pa for IDR^Dna2^ and IDR-3D^Dna2^ but not IDR-3A^Dna2^, consistent with Dna2 efficiently processing the substrate within the droplets. This observation was further supported by microscopy-based experiments showing significantly stronger decreases in Cy3-ssDNA signal within preassembled IDR^Dna2^ droplets upon the addition of full-length Dna2 (Supplementary Fig. 6f,g). In addition, at 30 mM NaCl, IDR-3D^Dna2^ formed the most stable droplets (Supplementary Fig. 6h,i) and more robustly supported function even when compared to IDR^Dna2^ (Supplementary Fig. 6j). Because all IDRs were purified from *E. coli*, the phosphomimetic substitutions act as fixed charge alterations, and the salt-dependent differences are therefore consistent with electrostatic tuning of condensate stability and material properties at different salt concentrations. Together, our results reveal that compared to IDR-3A^Dna2^, the IDR^Dna2^ and particularly the phosphomimetic IDR-3D^Dna2^ exhibit a stronger ability to operate in trans to boost Dna2 function within the in vitro conditions tested, even without adding factors such as RPA or Sgs1. In agreement with our findings, mutation of the three Cdk1-dependent phosphorylation sites examined here to alanine does not alter binding to Top3-Rmi1, MRX, or RPA complexes, nor the ability of RPA to stimulate resection in vitro^48^. This is consistent with phosphorylation modulating condensates rather than canonical protein-protein interactions.

To assess the impact of IDR status on untagged Dna2 in vivo, we CRISPR/Cas9-edited the wild-type *DNA2* open reading frame to generate the phosphomimetic Dna2-3D and phosphonull Dna2-3A versions of the full-length protein (Fig. 5c (top)). Using GFP-tagged versions of these endogenously expressed proteins, we first observed that Dna2 and Dna2-3D, but not Dna2-3A, efficiently formed foci following a one-hour zeocin treatment (Supplementary Fig. 7a,b). Considering that *EXO1* knockout prevents deleterious effects triggered by artificially modifying Dna2 phosphorylation sites^48^, we compared the effect of untagged Dna2, Dna2-3D, and Dna2-3A on resection, repair, and cell survival in an *exo1Δ* background. Following zeocin treatment, pRfa2 induction began early regardless of the full-length Dna2 variant expressed, resolving first for Dna2-3D and peaking last for Dna2-3A (Supplementary Fig. 7c). Concordantly, replacing wild-type Dna2 with Dna2-3D appeared to partly increase resistance to zeocin, but this did not reach statistical significance, while replacing Dna2 with Dna2-3A strongly and significantly increased sensitivity to zeocin (Fig. 5c (bottom); Supplementary Fig. 7d). We also assessed the ability of cells to resect and survive a single galactose-inducible and HR-reparable DSB on chromosome V and we confirmed that Dna2 foci rapidly form upon inducing the single DSB (Supplementary Fig. 7e,f)^61,62^. Cells with Dna2 or Dna2-3D displayed more efficient resection and repair of the single DSB compared to Dna2-3A cells (Fig. 5d; Supplementary Fig. 7g). These findings indicate that the Cdk1-dependent phosphorylation sites within the IDR of full-length Dna2 can boost phase separation and resection. Of note, our data indicate that the 3A mutation abolishes foci formation by full-length Dna2 in vivo and disrupts the material properties of droplets formed by the IDR alone in vitro. These observations suggest that phosphorylation of the Dna2 IDR both enables condensation under native physiological conditions and can tune condensate material properties that regulate enzymatic function.

### Evolutionary redistribution of condensate grammar

Since IDR^Dna2^ can boost catalysis even in trans, we examined this IDR more closely from an evolutionary perspective. To specifically assess how the condensate-forming capacity of Dna2 has evolved, we examined conservation of the protein’s disorder across eukaryotes. This analysis revealed that the IDR^Dna2^ disorder is present across fungal Dna2 proteins (Supplementary Fig. 8a-g) but absent from Dna2 orthologues in diverse non-fungal organisms, including humans (Supplementary Fig. 8a,h-s), consistent with our original computational analyses (Fig. 1c,d). Thus, the capacity for Dna2-encoded condensation appears to be largely restricted to fungi rather than a universally conserved property of the enzyme.

We next asked whether the molecular features of Dna2’s IDR are similar in different fungi. Comparative machine learning-based analyses showed that Dna2’s IDR in *S. cerevisiae* is more similar to that of the commensal pathogen *Candida albicans* than to that of *Aspergillus fumigatus*, *Schizosaccharomyces pombe*, or *Neurospora crassa* (Fig. 5e; Supplementary Fig. 9a-f). Concordantly, deletion of the IDR of *C. albicans* Dna2 markedly increased sensitivity to DNA damage (Fig. 5f; Supplementary Fig. 10a) and to the clinically used antifungal 5-fluorocytosine (Supplementary Fig. 10b), supporting a functional role for this fungal Dna2 IDR in genome maintenance and antifungal stress tolerance.

Finally, we asked whether the phase-separation grammar encoded by IDR^Dna2^ in yeast, but absent from human DNA2, has been lost or redistributed during evolution. Comparative analysis of candidate intrinsically disordered regions across human DNA repair proteins revealed patterns consistent with a redistribution of this organizational logic (Fig. 5g; Supplementary Fig. 10c (top)). The condensate-associated organizational features of *S. cerevisiae* IDR^Dna2^ showed strong similarity to predicted regions within human WRN, a RecQ helicase that promotes DNA2 function in human cells, as well as similarity to regions within the short-range resection regulators CtIP and NBN (Fig. 5g; Supplementary Fig. 10c (top)). These results are consistent with our earlier cluster analyses, in which we confirmed that Dna2 disorder features co-clustered with those of human WRN, NBN, and CtIP (Fig. 1d). These similarities do not necessarily imply functional equivalence but instead suggest shared organizational properties. That said, the IDR^Dna2^-overlapping region we identified in CtIP (amino acids 674-791; Fig. 5g) overlaps the previously defined 690-740 unstructured segment required for CtIP-dependent licensing of DNA2 function in human systems^63^, connecting the molecular grammar we identified to DNA2-regulatory function. Moreover, phosphomimetic substitution further increases the similarity between the disordered region of *S. cerevisiae* Dna2 and human CtIP (Supplementary Fig. 10c). Together, our observations are consistent with a model in which organizational features associated with enzyme-intrinsic condensation in fungi are redistributed across a restricted set of factors supporting DNA2-dependent resection in higher eukaryotes.

## Discussion

We propose that phosphorylation-tuned, Dna2-centered condensates promote long-range DNA end resection and thereby support checkpoint signaling, chromosome stability, and cell survival in *S. cerevisiae* (Supplementary Fig. 10d). Although these assemblies likely engage additional repair factors in vivo, our sufficiency, heterologous human IDR grafting, and phospho-tuning experiments demonstrate that Dna2 itself encodes a dominant and tunable condensation grammar that directly regulates its function. Consistent with this model, disruption of higher-order assembly impaired progression to advanced resection intermediates in vitro, linking condensation-dependent organization to sustained catalytic output. Collectively, our mechanistic studies reveal enzyme-intrinsic condensation as a previously unrecognized mechanism through which Dna2 activity can be regulated.

More broadly, our analyses suggest that condensation grammar can undergo evolutionary redistribution within conserved DNA repair pathways. In fungi, organizational capacity appears embedded within Dna2 itself, whereas higher eukaryotes redistribute this capacity across multiple repair-associated factors. The emergence of shared organizational features between fungal Dna2 and human resection proteins such as CtIP, WRN, and NBN is consistent with a more distributed organizational architecture underlying resection in higher eukaryotes. Such redistribution need not reflect direct transfer of sequence features between proteins but may instead emerge through progressive gain, loss, partitioning, or reinforcement of condensation-promoting features across interacting pathway components during pathway evolution. Whereas Dna2 in yeast couples catalytic activity with intrinsic condensate formation, higher eukaryotes may have distributed related organizational functions across multiple resection-associated factors. Such redistribution could allow conserved repair chemistry to be maintained while altering how higher-order organization is integrated into pathway regulation. Consistent with this possibility, the CtIP region identified here overlaps a previously defined unstructured segment required for human DNA2 function^63^, linking the organizational grammar identified in our study to resection in mammalian systems.

Our findings are also consistent with increasing evidence that spatial organization of DNA repair pathways can be externalized from catalytic machinery in mammals, including PARP1-dependent condensate signaling assemblies^32,33,35^ and MRE11-recruiting MRNIP condensates^28^. More broadly, species-specific deployment of condensate-associated features within conserved pathways is also reflected in the experimental confirmation of liquid-like properties of human RPA2 but not yeast Rfa2, as well as in the presence of extended unstructured regions within human CtIP that are absent from or significantly shortened in its yeast counterpart Sae2^21,24,29,63^. Our findings suggest that the redistribution of condensation-promoting grammar need not involve a simple transfer from enzymes to scaffolds. Rather, organizational features may be partitioned across, or reinforced within, distinct pathway components while preserving conserved biochemical outputs.

Our study also highlights several key questions and limitations. First, the grammar that promotes condensation, identified computationally within predicted intrinsically disordered regions, may in some cases be situated in dynamically structured or conditionally ordered segments. These segments can shift between disordered, oligomeric, or partially folded states depending on interaction partners, assembly conditions, or post-translational modifications. Additionally, the organizational features identified here might support a range of higher-order states, from liquid-like condensates to more transient or locally multivalent assemblies. Furthermore, although our comparative analyses suggest an evolutionary redistribution of condensation-promoting grammar across conserved repair pathways, understanding the exact evolutionary routes and selective pressures involved will require further expanding phylogenetic sampling and functional interrogation. Finally, regions in different human proteins show strong similarity to the fungal Dna2 condensate-associated features, but whether these human proteins deploy such grammar through bona fide condensates or other transient higher-order assemblies represents an important next step toward understanding spatial control of DNA repair in higher eukaryotes.

Collectively, our findings position phosphorylation-tuned, enzyme-intrinsic condensation as a regulatory principle of DNA end resection while supporting a model in which condensation grammar can be redistributed among components of conserved molecular pathways during evolution.

## Methods

### Yeast strain construction and research materials

Lithium acetate was used for all yeast transformations^62^. We validated endogenous tagging or deletions using colony PCR, growth in selection media, or microscopy, and point mutants and plasmids via sequencing. To generate the *dna2Δ* strain, a point mutation in the Pif1 locus (pif1-m2)^48^ was first introduced using Delitto Perfetto^64^ before a deletion cassette (*HPHMX6*) was used to replace the *DNA2* open reading frame. The Dna2-3A and Dna2-3D point mutations were CRISPR-Cas9-engineered^65^ within the endogenous *DNA2* locus. The lists of strains, antibodies, primers, sgRNA, labelled DNA substrates, and plasmids are included in Supplementary Tables 1-5. Regarding strain details and experimental setup, for Fig. 1e,f, and Supplementary Fig. 1h,i, the endogenous gene was directly GFP-tagged in the genome. For Fig. 1g-i, 4b,d, Supplementary Fig. 2c-f, and Supplementary Fig. 5b, GFP-tagged wild-type or variants of Dna2 were expressed under the control of the endogenous promoter from a centromeric plasmid. Strains used in Fig. 4c,e, and Supplementary Fig. 5a,c-h, were in a *pif1-m2* background to avoid *dna2Δ* lethality. For experiments comparing GFP-tagged, full-length wild-type Dna2, Dna2-3D, and Dna2-3A in Fig. 5c,d, and Supplementary Fig. 7a-d,g, the proteins were expressed from the endogenous *DNA2* locus, where the point-mutant variants were created using CRISPR-Cas9. For all genotoxic agent-induced DNA damage experiments, cells were treated with 0.03% (w/v) MMS (Cat. Nb. 156890050, Thermo Scientific) or 50 μg.ml^-1^ zeocin (Cat. Nb. R2500, ThermoFisher Scientific) for 1 hr at 30°C, except for the long-term cell survival spotting assays, which were conducted as described below.

### Plasmid construction

For *S. cerevisiae* studies involving plasmid-based expression of Dna2, a centromeric plasmid was used to express Dna2-GFP under its endogenous promoter^48^. The IDR-less Dna2Δ405-GFP plasmid was generated by deleting the first 405 amino acids while inserting an SV40 Nuclear Localization Signal (NLS) via site-directed mutagenesis^66^. To generate the IDR^FUS^-Dna2Δ405-GFP plasmid, the N-terminal IDR^FUS^ at residues 2-190 of FUS was PCR-amplified with flanking NheI and NarI restriction cut sites. Identical cut sites were introduced between the SV40 NLS and Dna2Δ405-GFP, allowing for the restriction enzyme-based digestion and annealing of the IDR^FUS^ onto the Dna2Δ405-GFP plasmid. Plasmids used to generate the CRISPR knock-ins^65^ were modified to generate gRNAs specific to the *DNA2* mutation target sites. For *C. albicans* studies, the promoter and coding sequence of *DNA2(A)* (A allele) or *DNA2(B)* (B allele) was amplified from SC5314 genomic DNA and cloned into a *pSFS2-SAT1* vector^67^, which had been modified to contain the *DNA2* downstream region between the XhoI/ApoI sites. The IDR (Ser2-Arg344) was truncated by fusion PCR and cloned into the same vector.

### Immunoblotting

Cells from logarithmically growing cultures (0.75 OD_600_) were treated with 0.2 M NaOH for 15 min on ice. Cells were pelleted, resuspended in Laemmli loading buffer, and boiled for 12 min at 95°C. Samples were centrifuged and the supernatants were loaded onto SDS PAGE gels for protein separation. Proteins were transferred onto nitrocellulose membranes and blocked in either 5% BSA (w/v) or milk in TBST (w/v) for 1 hr at 23°C (room temperature). Membranes were incubated in primary antibody at 4°C overnight, washed thrice in TBST (TBS with 0.1% Tween v/v), and incubated with HRP-conjugated secondary antibodies for 1 hr at 23°C. After three TBST washes, membranes were rinsed with Western ECL Substrate (BioRad) for 2 min before being developed using the ChemiDoc imaging system (BioRad).

### Microscopy

Cells from logarithmically growing cultures in synthetic complete media were treated with 0.03% w/v MMS (Cat. Nb. 156890050, Thermo Scientific) or 50 μg.ml^-1^ zeocin (Cat. Nb. R2500, ThermoFisher Scientific) for 1 hr at 30°C to induce DNA damage before the cells were pelleted and resuspended in 2 μg.ml^-1^ digitonin (Cat. Nb. D141-100MG, Sigma) alone or with 5% w/v 1,6-hexanediol (Cat. Nb. 240117, Sigma-Aldrich). Cells were then washed twice with 0.1 M potassium phosphate buffer before fixation with 3.7% (v/v) formaldehyde at 23°C for 5 min. The fixed cells were washed twice and resuspended in 100 μl of 0.1 M potassium phosphate buffer. For experiments involving nuclei staining, fixed cells were then incubated in 1μg.ml^-1^ DAPI (Cat. Nb. D1306, ThermoFisher Scientific) for 2 min before the wash steps. Non-damage-induced control cells were treated identically but in the absence of MMS or Zeocin. Fixed-cell images with Z stacks at 0.3 μm intervals were acquired with a Nikon C2+ confocal microscope (Nikon) using a 100X TIRF oil objective and a numerical aperture (NA) of 1.45. For imaging live cells expressing Dna2-GFP and Nup49-mCherry, cells subjected to DNA damage for 1 hr at 30°C were pelleted, washed with ddH_2_O, and resuspended in 100 μl of media. Cells were then imaged on a Leica SP8 confocal STED microscope (Leica) using a 100X oil objective with an NA of 1.4. GFP and mCherry excitation wavelengths corresponded to 488 nm and 584 nm, respectively, and emission spectra were captured using HyD detectors. Z-stacks of 0.3 μm sections were acquired using time-lapse microscopy to enable the reconstruction of foci behaviour in three dimensions over time.

### Droplet formation in vitro

In vitro droplet formation assays were performed on a parafilm-sealed 35-mm glass-bottom dish (Mattek) upon the addition of the eGFP-IDR^Dna2^ to an in vitro buffer-A (100 mM Tris-HCl, pH 8.0, 1 mM EDTA, pH 8.0, 2.5-5% PEG (PEG3350; w/v), and 25-400 mM NaCl as indicated in the figures). Using a Nikon C2+ confocal microscope (1.45 NA, 100X objective), droplet formation was visualised after 488 nm wavelength excitation and phase contrast imaging. Single-plane images were captured at the midplane of droplets over time.

### Nucleic acid-IDR condensate colocalization

First, equimolar amounts of Cy3-labeled ssDNA (Cy3-ssDNA) and IDR^Dna2^ (25 μM) were mixed and incubated for 5 min in the in vitro buffer-A (100 mM Tris-HCl, pH 8.0, 1 mM EDTA, pH 8.0, 2.5-5% PEG (PEG3350; w/v), and 100 mM NaCl) before being transferred to a 35-mm circular dish for imaging. Colocalization was measured by visualizing the eGFP-tagged IDR^Dna2^ droplets upon 488 nm wavelength excitation, and the Cy3-ssDNA upon 543 nm wavelength excitation. Successive image acquisitions for each channel were performed to prevent artifacts. Second, for *S. cerevisiae* yeast RPA (yRPA) experiments, 10 μM yRPA was added to the in vitro buffer-A (100 mM Tris-HCl pH 8.0, 1 mM EDTA pH 8.0, 2.5-5% PEG (PEG3350; w/v), and 100 mM NaCl) containing 25 μM IDR^Dna2^ and incubated for 15 min at 23°C, followed by the addition of Cy3-ssDNA to a final concentration of 10 μM. Imaging was then performed as stated above for the in vitro droplet formation assays. Third, colocalization of the Cy5-labelled DNA substrate (Cy5-DNA) with the IDR^Dna2^ droplets was visualised when 1 μM of Cy5-DNA was spiked into in vitro buffer-B (50 mM sodium phosphate, pH 7.0, 5% PEG (w/v), and 50 mM NaCl) with preformed IDR^Dna2^ droplets (25 μM). The Cy5-DNA and eGFP-tagged IDR^Dna2^ were imaged concurrently after excitation at 650 nm and 488 nm, respectively.

### Fluorescence recovery after photobleaching

Damage-induced Dna2-GFP foci and GFP-Tub1 were imaged similarly to the live-cell method above, with alteration. Only the GFP emission spectra on a single plane were captured with a PMT detector upon 488 nm wavelength excitation at 15% laser power. Three images were captured before the Region of Interest (ROI) containing the focus/filament was quenched with 100% laser intensity, followed by imaging of fluorescence recovery over 35 images acquired at 0.65 sec intervals. In vitro droplets were imaged and bleached on a Leica Stellaris 5 confocal microscope (Leica) equipped with a 1.3 NA 63X glycerol objective. Droplets were visualised after 0.2% laser excitation with 488 nm wavelength using a HyD detector. Four images were captured before a subsection of the droplet was bleached twice at 100% laser intensity, and 40 images were acquired at 1.29 sec intervals to monitor fluorescence recovery.

### Protein purification

*S. cerevisiae* Dna2 protein variants (full-length and Δ405) were expressed in the WDH886 strain^68^ using the pGAL-FLAG-HA-Dna2-His_6_ plasmid^69,70^ and pGAL-FLAG-HA-Dna2Δ405-His_6_ plasmid^71^, respectively. Dna2 proteins were purified as described previously^69^. The expression and purification of *S. cerevisiae* RPA have been described previously^21^.

His_6_-SUMO-eGFP-IDR^Dna2^ (wild-type, phosphomimetic, and phosphonull; codon optimized for expression in *Escherichia coli*) were expressed in BL21 (DE3) RIL cells transformed with pET-His_6_-SUMO-eGFP-IDR^Dna2^ plasmids encoding IDR^Dna2^, IDR-3D^Dna2^, and IDR-3A^Dna2^, respectively. Cultures were grown in TB containing 50 µg.ml^-1^ kanamycin and 25 µg.ml^-1^ chloramphenicol at 37°C with shaking until OD_600_ ∼ 0.6, at which time the temperature was reduced to 18°C. When the OD_600_ reached 0.8-1.0, cultures were induced with 1 mM IPTG for 18-20 h (18°C). Cells were harvested by centrifugation, washed once with ice-cold PBS, flash-frozen in liquid nitrogen and stored at -80°C until purification.

Frozen cell pellets (8-10 g) were thawed and re-suspended in 100 ml ice-cold lysis buffer (50 mM Tris-HCl pH 8.0, 400 mM NaCl, 10% (v/v) glycerol, 10 mM imidazole) supplemented with fresh lysozyme (0.1 mg.ml^-1^), protease inhibitors (0.1 mM PMSF, 10 µg.ml^-1^ aprotinin, 10 µg.ml^-1^ leupeptin, 10 µg.ml^-1^ pepstatin, 4 µg.ml^-1^ bestatin), and reducing agent (3 mM β-mercaptoethanol). Cells were lysed by sonication on ice using a Branson 250 digital sonifier (10-15 cycles of 30 sec on, 1 min or, 20% duty output). The extract was clarified by ultracentrifugation in a Beckman Coulter Optima LE-80K Ultracentrifuge with the Type 45 Ti rotor for 1 h 15 min at 142,000 *g* (4°C).

The supernatant was loaded onto an equilibrated 5 ml or 10 ml HisTRAP FF column (Cytiva), connected to an ÄKTA Pure25 FPLC (Cytiva). After loading, the column was washed with 25 column volumes (CV) of HisTRAP binding buffer (50 mM Tris-HCl pH 8.0, 400 mM NaCl, 10% (v/v) glycerol, 10 mM imidazole). Proteins were eluted in 1 CV fractions using a 10 CV linear gradient from 0%-75% HisTRAP elution buffer (50 mM Tris-HCl pH 8.0, 400 mM NaCl, 10% (v/v) glycerol, 500 mM imidazole), followed by a 5 CV step at 100% HisTRAP elution buffer. Peak fractions containing His_6_-SUMO-eGFP-IDR^Dna2^ were pooled and supplemented with in-house purified His_6_-Ulp1-His_6_ SUMO protease (added at approximately 1:50 mass ratio). The cleavage reaction was transferred to 15 ml Slide-A-Lyzer Cassettes with a 3.5 kDa MWCO (Pierce) and dialyzed against 4 L of HisTRAP binding buffer (overnight at 4°C).

After dialysis, His_6_-Ulp1-His_6_ and the cleaved His_6_-SUMO tag were removed by re-loading onto the HisTRAP FF column (Cytiva), which was equilibrated in HisTRAP binding buffer. The column was washed with a 10 CV step gradient from 0%-4% HisTRAP elution buffer, collected in 1 CV fractions. The eGFP-IDR^Dna2^ proteins were retrieved in the flow-through and early wash fractions, as confirmed by SDS-PAGE.

Fractions containing eGFP-IDR^Dna2^ proteins were pooled and diluted with TEG0 buffer (20 mM Tris-Cl pH 8.0, 10% (v/v) glycerol, 1 mM EDTA, 1 mM DTT) to adjust to ∼ 50 mM NaCl. The sample was further purified using a 5 ml HiTRAP Q HP column (Cytiva), equilibrated in HiTRAP Q binding buffer (20 mM Tris-Cl pH 8.0, 50 mM NaCl, 10% (v/v) glycerol, 1 mM EDTA, 1 mM DTT). After washing with 25 CV HiTRAP Q binding buffer, the column was developed with a series of linear and step gradients using HiTRAP Q binding buffer and HiTRAP Q elution buffer (20 mM Tris-HCl pH 8.0, 1 M NaCl, 10% (v/v) glycerol, 1 mM EDTA, 1 mM DTT). Fractions (2 ml) were collected in a 96-well plate (Eppendorf). This protocol was necessary to achieve maximum separation between eGFP-IDR^Dna2^ and various impurities.

The first step was a 1 CV linear gradient from 0%-10% HiTRAP Q elution buffer. The second was a step gradient using 10% HiTRAP Q elution buffer (10 CV), followed by another step gradient at 13% HiTRAP Q elution buffer (10 CV). The fourth step was a 10 CV linear gradient from 13%-20% HiTRAP Q elution buffer, followed by a 5 CV step gradient at 20% HiTRAP Q elution buffer. The final steps involved a 10 CV linear gradient from 20%-100% HiTRAP Q elution buffer, followed by a 5 CV step gradient at 100% HiTRAP Q buffer. Highly pure eGFP-IDR^Dna2^ was observed in fractions eluted with 13%-20% HiTRAP Q elution buffer, as determined by SDS-PAGE.

Peak fractions containing pure eGFP-IDR^Dna2^ were pooled and diluted 5-fold with NaP0 (50 mM sodium phosphate pH 7.0, 4 M NaCl) for hydrophobic interaction chromatography. The sample was loaded onto a 5 ml HiTRAP Phenyl HP column (Cytiva), equilibrated in HiTRAP Phenyl binding buffer (50 mM sodium phosphate pH 7.0, 4 M NaCl), and washed with 20 CV at 10% HiTRAP Phenyl elution buffer (50 mM sodium phosphate pH 7.0). Proteins were eluted in 1 CV fractions as follows: 10 CV step gradient from 10%-15% HiTRAP Phenyl elution buffer, 15 CV linear gradient from 15%-100% HiTRAP Phenyl elution buffer, and 5 CV step at 100% HiTRAP Phenyl elution buffer.

Fractions containing highly purified eGFP-IDR^Dna2^ proteins were identified by SDS-PAGE, pooled, and concentrated to approx. 12 mg.ml^-1^ (or 200 μM) in storage buffer (50 mM sodium phosphate pH 7.0, 200 mM NaCl) using Amicon 10 kDa MWCO Ultra-15 Centrifugal Filters (MilliporeSigma). Purified proteins were quantified spectroscopically (A_280_), aliquoted into protein lo-bind microfuge tubes, flash frozen in liquid nitrogen, and stored at -80°C. Prior to some *in vitro* droplet formation and nuclease activity assays, eGFP-IDR^Dna2^ proteins (wild-type, phosphomimetic, and phosphonull) were thawed and dialyzed against 4 L storage buffer (50 mM sodium phosphate pH 7.0, 200 mM NaCl) in Pur-A-Lyzer 1 kDa MWCO Midi Dialysis Tubes (MilliporeSigma) overnight at 4°C. The next day, samples were concentrated to 14-19 mg.ml^-1^ using Amicon 10 kDa MWCO Ultra-0.5 Centrifugal Filters (MilliporeSigma). Purified proteins were quantified spectroscopically (A_280_), aliquoted into protein lo-bind microfuge tubes, flash frozen in liquid nitrogen, and stored at -80°C.

### In vitro catalysis assays

Cy5-labelled 5’-flap DNA was generated by annealing oligonucleotides (Oligo1, Oligo2, and Oligo3) using previously established protocols^72^. Assays evaluating the impact of either 5% PEG-3350 (w/v)/5% 1,6-hexanediol (w/v) or of 25 μM eGFP-IDR/5% PEG-3350 were carried out as follows. First, reaction mixtures containing base buffer components (50 mM sodium phosphate pH 7.7, 30 mM or 50 mM NaCl, 2 mM MgCl_2_, 1 mM DTT, and 0.1 mg.ml^-1^ rAlbumin (New England Biolabs)) were prepared at room temperature. Next, the reactions were supplemented with variable buffer components (30% PEG-3350, 30% 1,6-hexanediol, or 200 μM eGFP-IDR^Dna2^ stock solutions), achieving final concentrations of either 5% PEG-3350, 5% 1,6-hexanediol, 5% 1,6-hexanediol/5%PEG-3350, 25 μM eGFP-IDR, or 25 μM eGFP-IDR/5% PEG-3350. The complete reaction mixtures were incubated at room temperature for 2-3 min, followed by the addition of 10X Cy5-labelled 5’-flap DNA (10 nM final) and subsequent equilibration for 5 min at 30°C. Processing was initiated by the addition of Dna2 or dna2Δ405 enzyme (1.1 nM final). Aliquots were taken at various time points and terminated by incubation with stop buffer (5X: 10 mg.ml^-1^ proteinase K, 10 mM CaCl_2_, 0.5% SDS (v/v)) at 37°C for 30 min. Stopped reactions were supplemented with DNA loading dye (6X = 30% (v/v) glycerol, 10 mM EDTA, 0.25% (w/v) Orange-G) and run on 10% native polyacrylamide gels for 75 min at 150 V. Gels were imaged on a Typhoon FLA 9500 instrument with a PMT voltage of 600V using an LPR (R665) filter (Cytiva). Quantification was conducted as described previously^54^. To visualize Dna2 nuclease activity within eGFP-IDR^Dna2^ droplets, 10 μM IDR^Dna2^ was incubated in resection assay buffer (50 mM sodium phosphate pH 7.7, 50 mM NaCl, 2 mM MgCl_2_, 1 mM DTT, 0.1 mg.ml^-1^ BSA, and 5% PEG) for 10 min to form the droplets before the addition of 100 nM Cy3-labelled ssDNA and incubation for 5 min. Upon confirmation of Cy3 signal colocalization within eGFP-IDR droplets, full-length Dna2 was spiked in at a final concentration of 4 nM, and images were captured at 1 and 2.5 min. The eGFP-IDR and Cy3-ssDNA were visualized following excitation at 488 nm and 543 nm, respectively.

### Microscopy image analysis

Analysis of fixed-cell and live-cell microscopy images, including quantification of standard fluorescence intensity, FRAP, and droplet measurements, was performed on Fiji (version 2.16), NIS-Elements AR software (version 4.10.00), or Imaris Viewer (version 10.0.1). Linear brightness and contrast adjustments, and a Gaussian filter, were applied equally across images within the same experimental condition. Raw images from the live-cell microscopy were analysed using Fiji (version 2.16). Fusion, fission, and dripping were scored following the confirmation of the tracked behavior via examination in 2D planes and 3D-reconstructed bodies. Live-cell images were also deconvolved using the Huygens Light Sheet Deconvolution Software before being viewed and processed on Imaris Viewer (version 10.0.1) to reconfirm the examined liquid-like behaviors. Linear pixel intensity was optimized in Imaris using the display adjustment tool.

### *S. cerevisiae* growth and sensitivity assays

10^5^ cells were spotted on the first column, followed by the spotting of 10-fold serial dilutions on plates of YEPD media alone, with 4 μg.ml^-1^ zeocin, or 0.01% (w/v) MMS for 4 days for initial screens, and with 10 μg.ml^-1^ zeocin or 0.02% (w/v) MMS for 5 days for subsequent characterizations. All spotting assays included sensitivity controls such as Rad52-deficient or Dna2-deficient cells.

### Single DNA double-strand break assays

The TGI354 system was used to measure cell survival following induction of an HO-inducible genomic DNA double-strand break (DSB)^61,62^ on chromosome V, repairable via an incomplete *MAT* locus (*MATinc*) template on chromosome III. Logarithmically growing cells in YEP with 2% raffinose (v/v) were plated on YEP plates containing 2% glucose (v/v) or 2% galactose (v/v). Cell survival was calculated by dividing the colony-forming units (CFU) from YEP with 2% galactose plates by the CFUs from YEP with 2% glucose plates.

### DNA end resection in vivo

Resection was measured per an established protocol^73,74^ with modifications. Cells with the engineered HO break site on chromosome V (TGI354)^61,62^ were grown in YEP with 2% raffinose (v/v) to log phase before being transferred to YEP with 2% raffinose (v/v) and 2% galactose (v/v) media to induce the HO cut. Roughly, 10^7^ cells were collected at different time points and treated with DNA extraction buffer (50mM Tris-HCl pH 8.0, 1% SDS (w/v), 100 mM NaCl,10 mM EDTA, 1% beta-mercaptoethanol (v/v), and 20 units of zymolyase 20T (Cat. Nb. ZYM001.250, Bioshop) per extraction for 10 min at 37°C with shaking at 300 rpm. DNA was then precipitated after phenol-chloroform extraction and resuspended in 100 μl ultrapure water. Samples were split into two, where half the sample was digested with EcoRI (Cat. Nb. R3103, New England Biolabs) in rCutSmart buffer (Cat. Nb. B6004, New England Biolabs) for 1 hr at 37°C. 2 μl of the digested or undigested DNA (0.2 ng μl^-1^), 5 μl of SyBr (BIO-98050, Bioline), 0.2 μl of each primer and 2.6 μl of ultrapure water were added to each well before qPCR was performed with the following settings: 1 time (95°C for 1 min) and 39 times (95°C for 15 sec then 57°C for 20 sec). The qPCR was performed with three technical replicates per sample per biological replicate, and *β*-actin was used as an internal control. The percentage of ssDNA was calculated as described^73,74^.

### Contour-clamped homogeneous electric field electrophoresis

Logarithmically growing cells (OD_600_ = 1.7) were collected, washed twice with 0.05 M EDTA, and processed as described^75–77^ with modifications. Cells were resuspended in 43.5 μl of 0.05 M EDTA along with 1.5 μl zymolyase 20T (10 mM sodium phosphate, 50% (v/v) glycerol, 10 mg ml^-1^). 45 μl of prewarmed, low-melting agarose (1.2% (w/v) in 0.125 M EDTA) was then added and solidified in plug molds (Cat. Nb. 1703713, BioRad) at 4°C. Gel plugs were then incubated in 7.5% beta-mercaptoethanol (v/v) and 0.5M EDTA at 37°C overnight. This was followed by another incubation in NDSK buffer (0.5 M EDTA, 10 mg ml^-1^ N-lauryl sarcosine, 1mg ml^-1^ proteinase K) at 50°C overnight. Gel plugs were then run on a Contour-clamped homogeneous field electrophoresis (CHEF)-specific agarose gel (1% w/v) and resolved in 1X TAE using a CHEF DR II apparatus (BioRad). After 48 hr of electrophoresis (3 V cm^-1^, 0.2-266 sec), the gel was stained with ethidium bromide and imaged.

### *C. albicans* strains and dose-response assays

*DNA2* was deleted using transient CRISPR technology^78^ in *C. albicans* reference strain SC5314. The sgRNA was generated by gene-specific primers oLC14458/14459 and the repair cassette was amplified by oLC14460/14461 from XmnI linearized *CaNAT-FLP* pLC49 vector^79^. Nourseothricin (150 μg.ml^-1^)-resistant transformants were genotyped using oLC14462/14463 to confirm the absence of the *DNA2* coding region and then grown in YCB-BSA medium (1.17% yeast carbon source supplemented with 0.2% BSA (w/v)) to excise the NAT marker. One representative mutant was selected, and then full-length or IDR-less *DNA2* alleles were reintroduced in tandem with a *SAT1* marker, which were liberated by SacI/ApaI digestion from the corresponding *pSFS2* plasmids (Supplementary Table 5). Integration was facilitated by transient CRISPR using sgRNA generated by oLC11150/11151, which targeted the ‘scar’ sequence after excision of the *NAT* marker used for gene disruption. Integration at both *dna2*-deleted loci was checked by loss of the ∼900-base-pair amplicon of oLC14469/14470. The *SAT1* marker was excised by culturing in YPmaltose (2%).

For dose response assays, all strains were freshly streaked from frozen stocks and maintained at room temperature on YPD agar for no more than one week before experimentation. Dose-response assays were performed as described^80^. The inoculum was diluted from a saturated overnight YPD culture and added to two-fold compound gradients generated by manual pipetting or direct dispensing with a Tecan D300e liquid dispenser. The highest concentrations tested were 0.1% (v/v) for MMS (Sigma-Aldrich), 1 mg.ml^-1^ for Zeocin, and 0.2 μg.ml^-1^ for 5-fluorocytosine (5-FC; Sigma-Aldrich). Except for 5-FC, all compounds were tested in YPD, and growth was measured by OD_600_ after a 24-hour incubation at 30°C. For 5-FC, growth was measured in synthetic-defined medium (0.67% yeast nitrogen base; 2% glucose) after 48 hours. All tests were performed in technical duplicates and repeated in independent experiments or using multiple transformants. Results were shown in heatmap format, with the growth of the wild-type strain in compound-free medium set to 1.

### Bioinformatics

Prediction of disordered protein content across the 312 DDR proteins and 5,737 control non-DDR proteins was conducted using IUPred and PONDR-VSL2 tools^40–42^. Protein sequences were acquired from UniProt (available at uniport.org) and filtered for reviewed sequences (Swiss-Prot). Disordered domains were acquired using the IDR prediction software IUPred (available at iupred1.elte.hu) and VSL2 (available at pondr.com)^41^. Monomeric protein disorder predictions were conducted without filtering out conditionally structured domains to avoid any biasing of the analyses. Prediction of *S. cerevisiae* Dna2 protein structure, including high and low confidence regions was determined using AlphaFold^81^.

For the analysis of IDR grammar, intrinsically disordered regions (IDRs) with a minimum length of 30 residues were identified using the VSL-2 predictor^41^ and the MobiDB database (version 4.0)^82^. Collected IDR sequences were analyzed using the NARDINI+ algorithm to extract 90 molecular grammar features^44^. To validate the use of molecular grammars rather than primary-sequence identity alone, the collected IDRs were ranked using BLOSUM62, which did not recapitulate the structure observed with NARDINI+. For example, one of the top hits (NBN region 184-220) was ranked 56^th^ in similarity by sequence identity, confirming that NARDINI+ molecular grammars capture information not available from primary-sequence comparisons alone. NARDINI+ was also selected over a protein language model to accommodate the wide range of IDR sizes and enable cross-species comparison. Grammar heatmaps and *H. sapiens* cluster distances figures were generated using code available in NARDINI+. The Z-scores were calculated relative to a fixed reference proteome from *S. cerevisiae*. Z-score vectors were pre-processed (NaN handling, min variance cutoff (0.1%), and re-standardized using StandardScaler form sklearn.preprocessing). Therefore, the resulting clustering is relative to the chosen input IDRs, and clustering is done on the relative profile of features rather than on raw magnitude. For charge distribution analysis, IDRs were identified as described above and CIDER was used to calculate linear net charge per residue and generate charge distribution plots^83^. For UMAP visualization, the 90 features were renormalized to a fixed reference proteome from *H. sapiens*. Three PCs were fed into UMAP, and Hierarchical clustering was performed on the resulting manifold.

For feature selection via XGBoost and Clustering, an XGBRegressor model was trained on the Z-score vectors with sequence indices (0, 1, 2, …, N−1) as the target variable^84^. Feature importance scores were computed from the trained XGBoost model, and the top 15 ranked features were selected for 2D principal component analysis (PCA). Clustering quality was assessed using Silhouette score, Davies-Bouldin index, and Calinski-Harabasz index. For hyperparameter optimization and sensitivity analysis, 92 hyperparameter configurations were tested per dataset, with configurations of n_estimators (80-300) and max_depth (3-6). Silhouette score, Davies-Bouldin index, and Calinski-Harabasz index were computed. Final visualizations for each dataset were generated using hyperparameters that exhibited high Silhouette scores and resided within stable regions (Dna2 homologs: n_estimators=90, max_depth=3; Human repair proteins: n_estimators=150, max_depth=5). To confirm robustness across the search space, Spearman correlation and coefficient of variation (CV) of pairwise distance vectors across 92 hyperparameter configurations were computed across all datasets, confirming high consistency in the relative ordering of distances (mean Spearman ρ=0.712; 50.2% of proteins CV<0.25; 23.1% of proteins CV<0.10).

### Statistical analysis

General statistical analysis was performed in GraphPad Prism (version 10.6.1) using the indicated tests as best suited for the specific experimental design and type of data analysis.

Unless otherwise indicated, replicate information is as follows. The quantification data were generated using at least three independent biological or cell-free replicates. Microscopy images, gels, and blots are representative of at least three independent biological replicates. The total number of cells analyzed is indicated in the figure legends, where applicable. All statistical comparisons in the disorder grammar clustering analysis were two-tailed, and metrics were computed without assumptions regarding cluster separability or shape.

### Code

Following the use of NARDINI+ to compute molecular grammars as described in the methods, new code used for XGBRegressor-based feature selection and clustering analysis includes: <u>enhanced_clustering_exploration_dynamic.py</u> for testing top n clusters (exploratory, data-set dependent); <u>XGBoost_hyperparam_grid.py</u> for hyperparameter search and stability-metrics computations; <u>enhanced_cluster_visualizer_dynamic_S14.py</u> for implementing the complete analysis pipeline (data preprocessing, feature selection via XGBoost, and PCA clustering); <u>clustering_robustness_analysis.ipynb</u> for hyperparameter sensitivity validation across 92 configurations (n_estimators: 80-300, max_depth: 3-6). All code is written in Python 3.8+ and requires publicly available packages (pandas, NumPy, scikit-learn, SciPy, XGBoost, Matplotlib).

## Acknowledgements

We thank Drs. Petr Cejka, Grant W. Brown, and Grzegorz Ira for materials. We thank Drs. Matthias Altmeyer, Rahul G. Krishnan, Hyun O. Lee, Marc Meneghini, Alan M. Moses, Taekjip Ha, and Mekhail lab members for discussions. We thank Dr. Paul Paroutis at the SickKids Children’s Hospital Imaging Facility for technical advice. R.H. is supported by Canadian Institutes of Health Research (CIHR) grants (183597, 186072). L.E.C. is supported by a CIHR grant (FDN154288) and a Canada Research Chair (CRC, Tier 1) in Microbial Genomics & Infectious Disease and is co-Director of the CIFAR Fungal Kingdom Threats & Opportunities program. H.D.M.W. is supported by CIHR grants (156297, 487044) and a CRC in Mechanisms of Genome Instability (950231487, 202100346). This research was primarily supported by CIHR grants (180469, 190143) and a CRC in Spatial Genome Organization (950230661) to K.M., who is supported by the Royal Society of Canada.

## Author contributions

C.A.J. and K.M. conceived and designed the project and wrote the manuscript, which was edited by the other authors. C.A.J. and G.L. designed and conducted the initial in silico and in vivo disorder content proteome screens. C.A.J. designed, conducted, and analyzed most experiments, with assistance from J.N.Y.C to CRISPR-Cas9 yeast genome editing and T.C. to CHEF. J.T. and H.D.M.W. purified proteins used for in vitro experiments, and B.J.P. designed and conducted in vitro resections. J.L. designed and conducted comparative analyses of condensate grammar with guidance from C.A.J. and K.M., and assisted C.A.J. with in vivo resections. Z.L. designed and conducted in vivo *C. albicans* experiments. R.H. supervised T.C., L.E.C. supervised Z.L., H.D.M.W. supervised B.J.P. and J.T., and K.M. supervised C.A.J., G.L., J.L., J.N.Y.C., and T.C.

## Generative AI and AI-assisted technologies

During the writing process, the authors used Grammarly and OpenAI to review grammar and in some cases to improve readability of the main text. The authors carefully reviewed recommendations from these tools and are responsible for the content.

