## Supplementary Information for "Dna2-intrinsic condensation regulates DNA end resection and reveals evolutionary redistribution of condensate grammar"

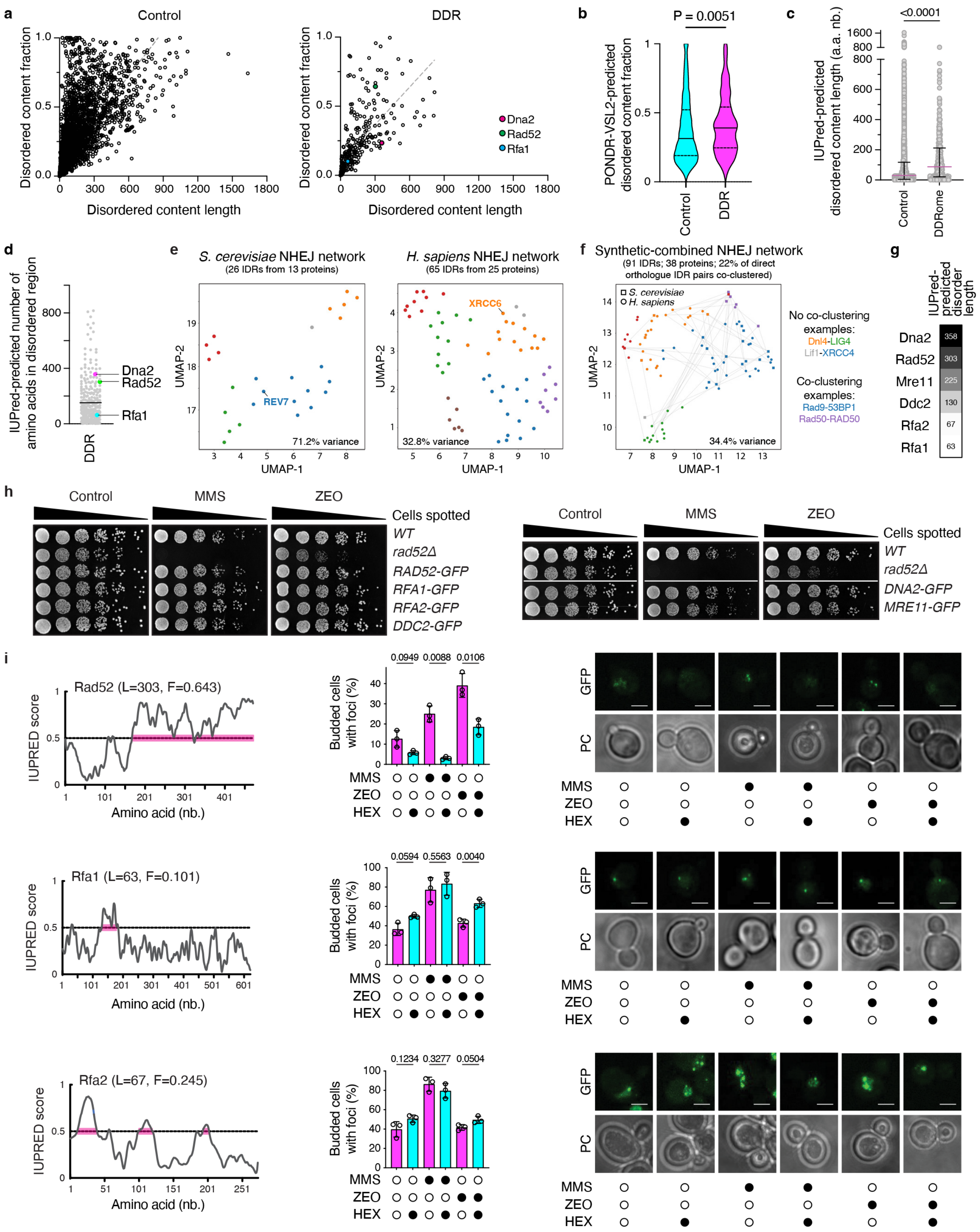


**Supplementary Fig. 1. Intrinsic disorder predictions and validation of DDR protein tagging and condensate sensitivity. a**, Distribution of *S. cerevisiae* DNA damage response (DDR) proteins and control non-DDR proteins by IUPred-predicted intrinsic disorder length and fraction of full-length protein. *n* = 5,737 control proteins and 312 DDR proteins; dashed line, simple linear regression. **b**, Disordered fraction of DDR proteins and control proteins predicted by PONDR-VSL2. Data are shown as the mean ± interquartile range; *n* = 5,737 control proteins and 312 DDR proteins; two-tailed unpaired t-test. **c**, IUPred-based disordered region length (right) in DDR and control proteins. Data are shown as the mean ± interquartile range; *n* = 5,737 control proteins and 312 DDR proteins; two-tailed Mann-Whitney test. **d**, Disordered region length distribution across the DDRome. The mean is indicated with a horizontal line. **e**-**f**, UMAP projections from machine-learning comparative analyses of condensate grammar of intrinsically disordered regions (IDRs) within non-homologous end-joining (NHEJ) proteins. Points represent individual IDRs labeled by hierarchical clustering assignment. Projections are separated by species (*S. cerevisiae*, left; *H. sapiens*, right) in (**e**), with lines connecting direct yeast-human orthologues in (**f**). Conserved pathway components with disorder in only one of the two species are indicated (**e**). **g**, DDR proteins with a range of disordered content length. **h**, Survival assay of *S. cerevisiae* cells harboring GFP-tagged DNA repair proteins on control media or on media supplemented with methyl methanesulfonate (MMS, 0.01%) or zeocin (ZEO, 4 $\mu$g.ml$^{-1}$) following a four-day incubation. Rad52 knockout cells served as a sensitivity control. **i**, IUPred-predicted disorder (left; with disorder length (L) and fraction (F) scores indicated), quantification of repair protein foci induced by MMS (0.03% w/v, 1 hr) or ZEO (50 $\mu$g.ml$^{-1}$, 1hr), their sensitivity to 1,6-hexanediol (HEX) (middle), and representative repair foci images (right). Quantification data are shown here as the mean ± s.d.; *n* = 3 independent biological replicates, two-tailed unpaired *t*-test with Welch's correction, and the means are also shown in the heatmap in Fig. 1e; scale bars, 3 $\mu$m.

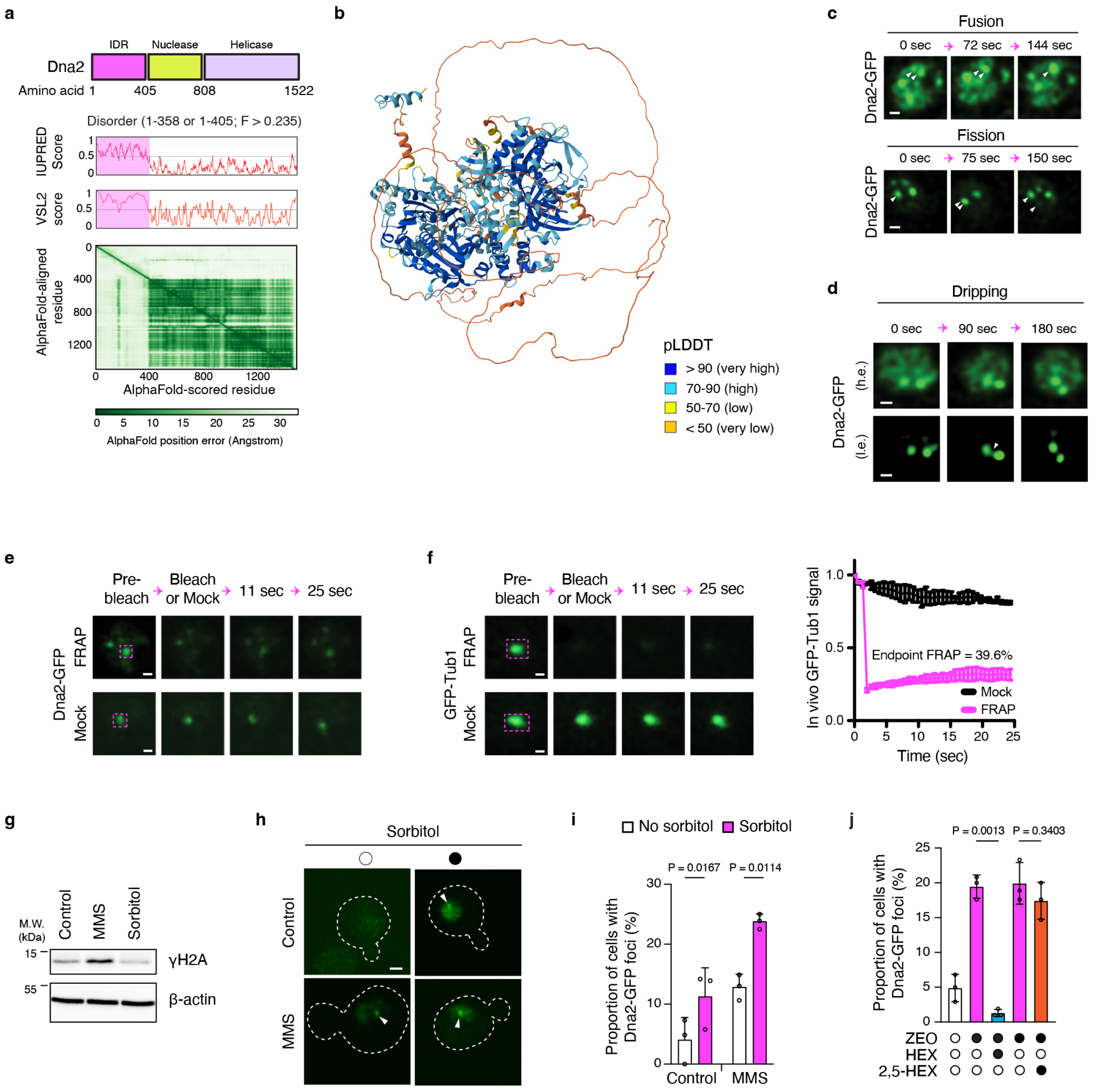


**Supplementary Fig. 2. Additional liquid-like behaviors and stress-induced assembly of Dna2 condensates in vivo. a**, Schematic of *S. cerevisiae* Dna2 (top), IUPred and PONDR-VSL2 disorder scores (middle), and AlphaFold positional error scores highlighting the putative intrinsically disordered region (IDR) spanning residues 1-405. **b**, AlphaFold-predicted structure of *S. cerevisiae* Dna2, highlighting the unstructured N-terminal region (residues 1-405, orange). pLDDT, predicted local distance difference test. **c**, Additional representative examples of fusion (with relaxation back to a spherical shape) and fission of Dna2-GFP repair foci following a 1 hr MMS treatment. Arrowheads highlight fusion (top) and fission (bottom). **d**, Additional high-exposure (h.e.) and low-exposure (l.e.) images of Dna2-GFP foci undergoing dripping (arrowhead), as shown in Fig. 1g. Scale bars, 0.5 μm. **e**, Representative images from fluorescence recovery after photobleaching (FRAP) analysis of Dna2-GFP repair foci in MMS-treated cells. Scale bars, 0.5 μm. **f**, Representative images (left) and quantification (right) from FRAP analysis of the negative control GFP-Tub1 in MMS-treated cells. Data show the mean ± s.d.; *n* = 30 (FRAP) or *n* = 23 (Mock) cells. **g**, Confirmation that sorbitol does not increase levels of the DNA damage marker γH2A. Cells treated with MMS served as a control. Scale bars, 0.5 μm. **h**,**i**, Representative images (**h**; scale bar, 1 μm) and quantification (**i**) showing that sorbitol induces Dna2-GFP foci formation (arrowheads) in untreated control cells and in MMS-treated cells. Data show the mean + s.d. from *n* = 3 independent biological replicates, two-tailed paired *t* test (**i**). **j**, Differential sensitivity of MMS-induced Dna2-GFP foci 1,6-hexanediol (HEX) versus the less active positional isomer 2,5-hexanediol (2,5-HEX). Data show the mean ± s.d.; *n* = 3 independent biological replicates, two-tailed unpaired t test with Welch's correction. The control, ZEO, and ZEO+HEX conditions are the same as in Fig. 1f, facilitating comparison.

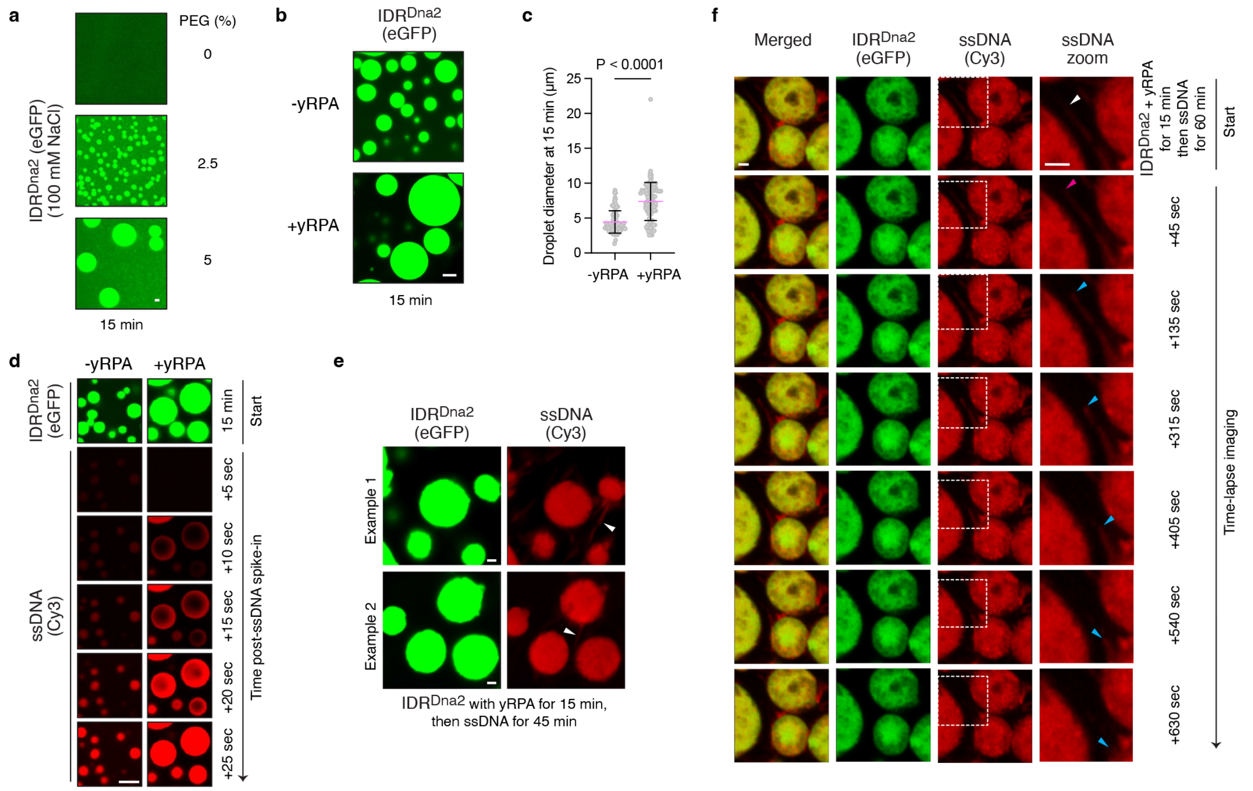


**Supplementary Fig. 3. Biochemical characterization of IDR$^{Dna2}$ and its intersection with yRPA and ssDNA. a**, Dependence of IDR$^{Dna2}$ droplet size on polyethylene glycol (PEG) concentration in vitro. **b**,**c**, Representative images (**b**) and quantification (**c**) showing the effect of the yeast RPA complex (yRPA) on IDR$^{Dna2}$ droplet size in vitro. Data show the mean ± s.d.; *n* = 86 (-yRPA) and 100 (+yRPA) droplets, two-tailed Mann-Whitney test (**c**). **d**, Effect of yRPA on the capacity of IDR$^{Dna2}$ droplets to attract ssDNA (10 µM). **e**,**f**, Snapshot images (**e**) and time-lapse microscopy (**f**) showing the reorganization of ssDNA by IDR$^{Dna2}$ assemblies in the presence of yRPA. Arrowheads denote ssDNA-positive filament presence (**e**), and their dynamics (**f**) are monitored at the starting point (white arrowhead), extension stage (magenta arrowhead), and shortening stage (cyan arrowhead). 25 µM IDR$^{Dna2}$, 5% PEG, 100 mM NaCl, and 10 µM yRPA was used (**b**-**f**). Scale bars, 1 µm (**a**, **e**, **f**) or 5 µm (**b**, **d**).

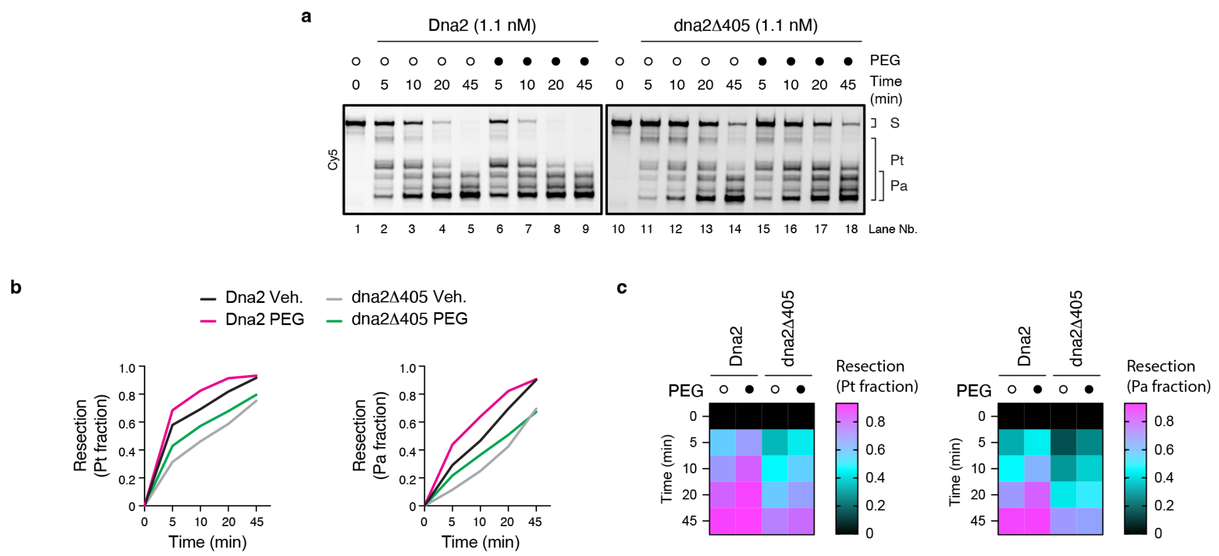


**Supplementary Fig. 4. Additional characterization of wild-type and IDR-less Dna2 nuclease activity kinetics in vitro.** Gels (**a**) and quantification (**b**-**c**) of full-length Dna2 or IDR-less Dna2 (dna2Δ405) activity based on the Pt fraction or Pa fraction in the presence or absence of 5% polyethylene glycol (PEG). Quantification results are displayed as line graphs (**b**) and heatmaps (**c**) to provide different perspectives of the same data.

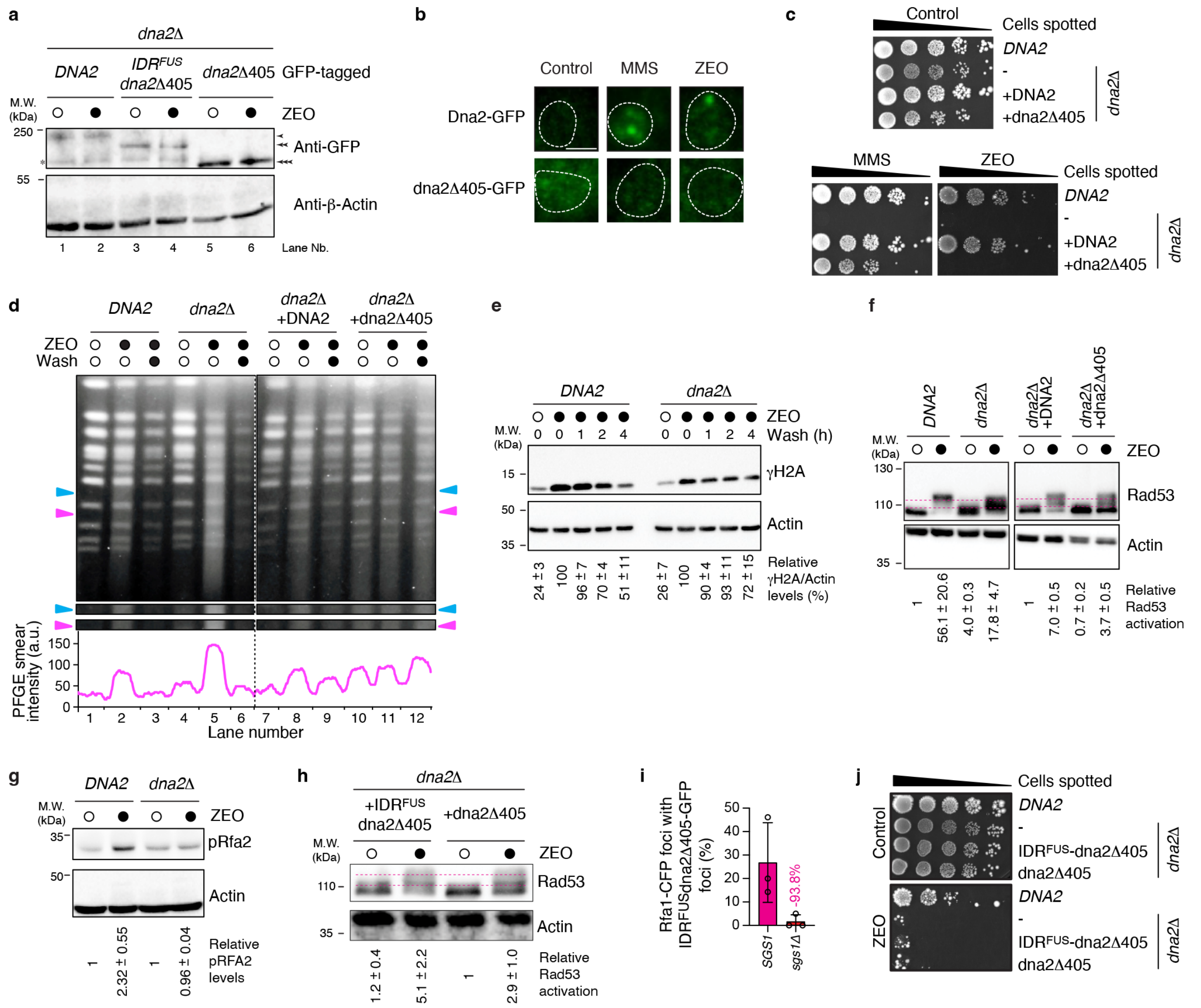


**Supplementary Fig. 5. Genetic studies of IDR$^{Dna2}$ function and IDR$^{FUS}$-mediated rescue.** **a**, Anti-GFP immunoblotting showing expression of full-length Dna2 and IDR-less Dna2 with or without fusion to IDR$^{FUS}$. Arrowheads indicate the expected sizes. **b**,**c**, Effect of IDR$^{Dna2}$ removal on GFP-tagged protein foci formation (**b**) and long-term cell survival (**c**) following treatment with methyl methanesulfonate (MMS) or zeocin (ZEO). Results are representative of 3 independent experiments. **d**, Contour-clamped homogeneous electric field (CHEF) electrophoresis showing the effect of losing Dna2 (lanes 1-6) or replacing it with plasmid-encoded full-length or IDR-less Dna2 (lanes 7-12) on genome stability following exposure to ZEO, and after its removal for two hours. Arrowheads indicate gel regions with DNA smearing, shown enlarged in gel strip insets and corresponding intensity trace plots. **e**, Effect of Dna2 loss on the resolution of ZEO-induced γH2A accumulation. Quantification shows mean ± s.e.m., *n* = 3 independent biological replicates. **f**, Effect of Dna2 loss (lanes 1-4) or its replacement with plasmid-encoded full-length or IDR-less Dna2 (lanes 5-8) on Rad53 checkpoint activation following treatment with ZEO. Activation scores reflect relative band upshifts and signal intensity. Quantification shows mean ± s.e.m., *n* = 3 independent biological replicates. **g**, Immunoblots and quantification showing the effect of Dna2 loss on zeocin-induced phosphorylated Rfa2 (pRfa2) levels. Quantification shows the mean ±

s.e.m.; *n* = 3 independent biological replicates. **h**, Effect of fusing IDR$^{FUS}$ to the IDR-less Dna2 on ZEO-induced Rad53 activation. Quantification shows mean ± s.e.m., *n* = 3 independent biological replicates. **i**, Assessment of the localization (mean ± s.d.) of IDR$^{FUS}$-dna2Δ405-eGFP foci to Rfa1-CFP-marked DNA damage sites in *SGS1* and *sgs1Δ* cells treated with ZEO. **j**, Assessment of long-term cell survival in the presence of ZEO. **a**-**j**, ZEO treatment was 50 μg.ml$^{-1}$ for 1 hr (**a**-**b**,**d**-**i**) or 10 μg.ml$^{-1}$ for 5 days (**c**) or 7 days (**j**), and MMS concentration was 0.03% for 1 hr (**b**) or 0.02% for 5 days (**c**).

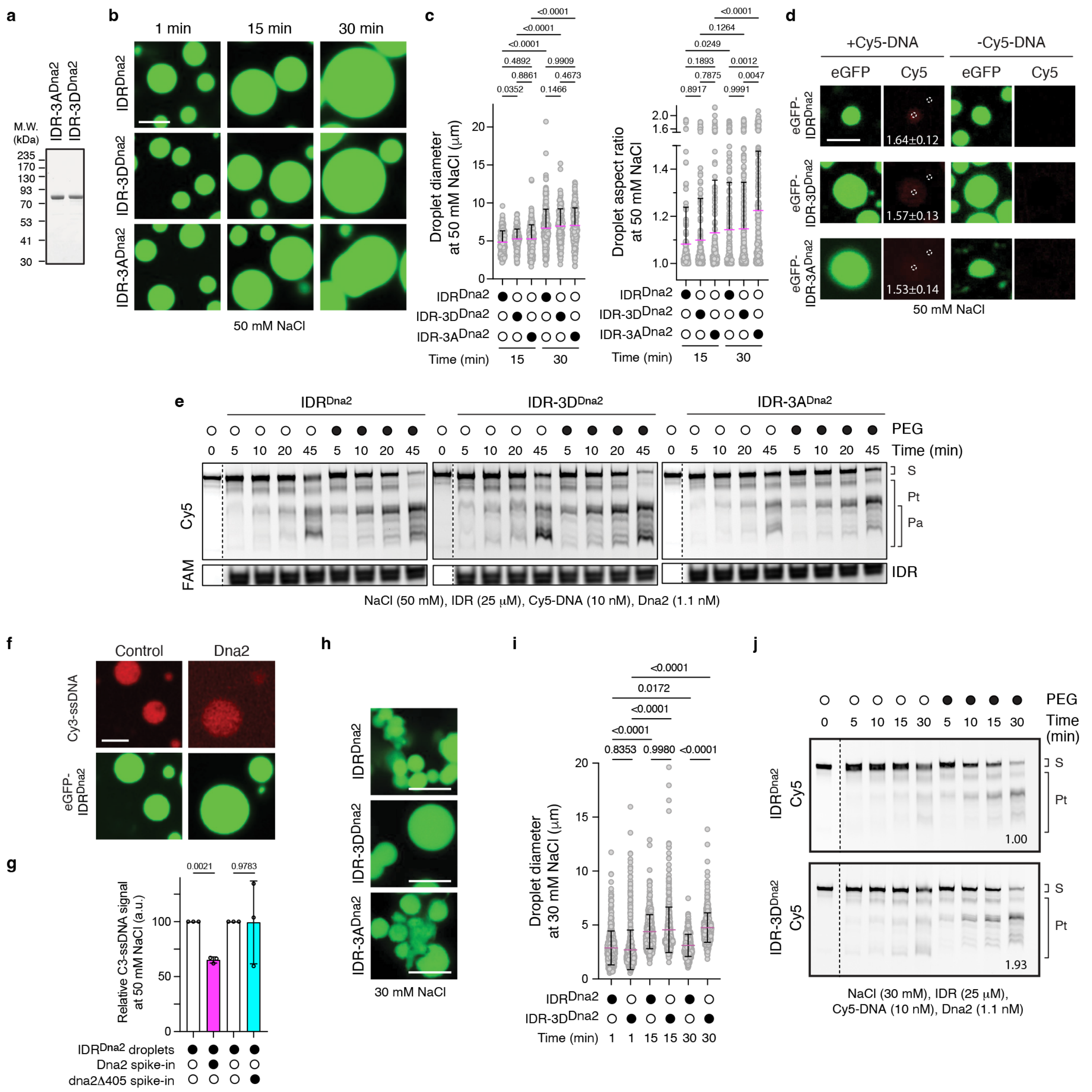


**Supplementary Fig. 6. Biophysical and function-modulating properties of phospho-tuned IDR$^{Dna2}$ droplets in vitro. a**, Purification of IDR$^{Dna2}$ variants that are either phosphonull (IDR-3A$^{Dna2}$) or phosphomimetic (IDR-3D$^{Dna2}$) from *E. coli*. **b**,**c**, Representative images (**b**) and quantification of droplet diameter (**c**, left) and aspect ratio (**c**, right) for droplets formed by IDR$^{Dna2}$, IDR-3D$^{Dna2}$, and IDR-3A$^{Dna2}$ at 50 mM NaCl. Scale bar, 5 μm. Data in **c** show the mean + s.d., 3 independent cell-free replicates, two-way ANOVA with Tukey's test; from left to right for droplet diameter, *n* = 226, 268, 207, 318, 154, and 191 droplets, and from left to right for droplet aspect ratio, *n* = 173, 200, 185, 196, 235, and 195. **d**, Representative images and quantification showing comparable recruitment of Cy5-labeled ssDNA into droplets formed by wild-type IDR$^{Dna2}$ (25 μM; 50 mM NaCl) and its phosphomimetic and phosphonull variants following a 15 min incubation. Quantification shows the ratio of Cy5 signal intensity inside the eGFP droplet compared to outside of the droplet. Quantification shows the mean ±

s.e.m.; *n* > 5 droplets per condition. Scale bar, 1 μm. **e**, Representative gels of in vitro nuclease activity assays testing the *in trans* effect of preformed $IDR^{Dna2}$ droplets or variants on full-length Dna2 function at 50 mM salt. The Cy5-labeled 5'-ssDNA flap substrate (S), total products (Pt), and advanced products (Pa) are indicated. Quantification from 3 independent replicates is in Fig. 5d. **f**, Representative images showing Cy3-ssDNA signal within in vitro preassembled $IDR^{Dna2}$ droplets decreases upon spiking in full-length Dna2. **g**, Quantification of C3-ssDNA signal within preassembled $IDR^{Dna2}$ droplets following a spike-in of full-length Dna2 or the IDR-less dna2Δ405. Data show the mean ± s.d.; *n* = 3 independent cell-free replicates each from at least 25 droplets, paired two-tailed *t* test. **h**,**i**, Representative images illustrating how IDR-$3D^{Dna2}$ forms droplets that are larger and more spherical than $IDR^{Dna2}$ or IDR-$3A^{Dna2}$ at 30 mM NaCl (**h**; scale bar, 5 μm) and detailed droplet size quantification (**i**) for $IDR^{Dna2}$ and IDR-$3D^{Dna2}$ showing the mean ± s.d.; from left to right, *n* = 646, 729, 526, 577, 522, 573 droplets, respectively, from 3 independent cell-free replicates, two-way *ANOVA* with Sidak's test. **j**, Results of an in vitro assay testing the *in trans* effect of preformed $IDR^{Dna2}$ and IDR-$3D^{Dna2}$ droplets on the activity of full-length Dna2 at 30 mM salt. The Pt fraction-based fold change relative to the wild-type $IDR^{Dna2}$ is indicated.

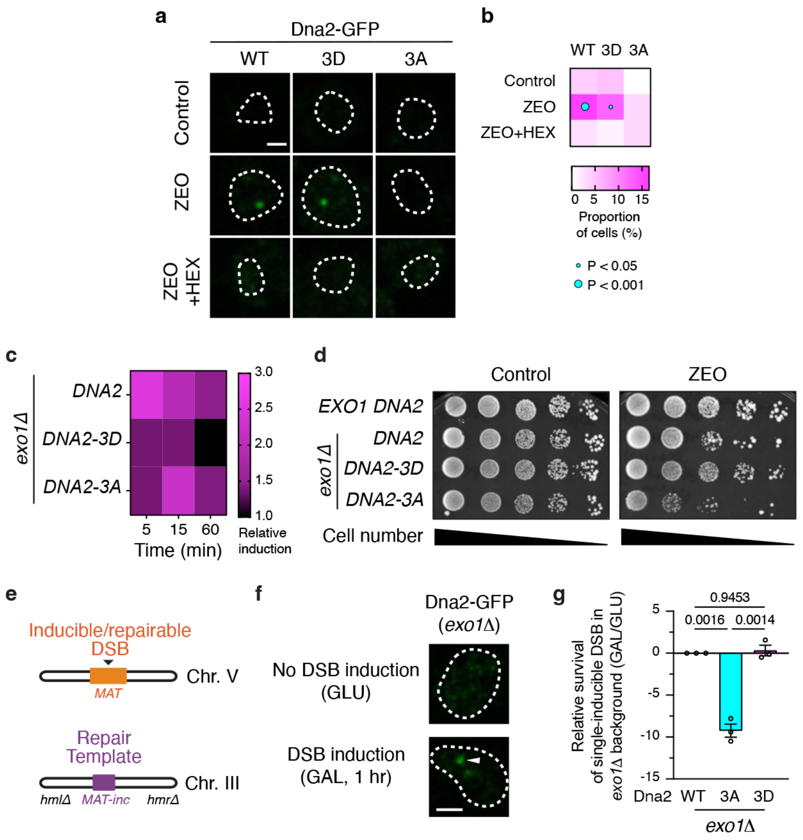


**Supplementary Fig. 7. Phosphoregulation of Dna2 condensation and function in vivo.** **a**,**b**, Representative images (**a**) and quantification (**b**) repair foci formation induced following a one-hour treatment with 50 μg.ml$^{-1}$ zeocin (ZEO) in cells expressing GFP-tagged Dna2, Dna2-3D, or Dna2-3A in the absence or presence of 1,6-hexanediol (HEX). Data (**b**) show the mean; from the heatmap's top left corner to the bottom right corner, $n$ = 120, 102, 89, 97, 121, 89, 104, 97, and 91 cells from 3 independent biological replicates, two-way ANOVA with Tukey's test. Scale bar, 1 μm. **c**, Effect of Dna2, Dna2-3D, and Dna2-3A on ZEO-induced Rfa2 phosphorylation (pRfa2) levels. **d**, Representative spotting assays showing the effect of full-length Dna2 and its phosphomimetic (Dna2-3D) and phosphonull (Dna2-3A) variants on cell survival in the presence of 10 μg.ml$^{-1}$ ZEO for 5 days. Quantification is shown in Fig. 5c. **e**, Schematic of the single inducible chromosome V DSB repaired by homologous recombination using donor sequences on chromosome III. **f**, Expression of Dna2-GFP under control of the endogenous *DNA2* promoter from a plasmid confirms repair foci form within 1 hr of inducing the single DSB. Scale bar, 1 μm. **g**, Effect of Dna2, Dna2-3D, and Dna2-3A on the survival of the single inducible DSB. Data are shown as mean ± s.e.m., $n$ = 3 independent biological replicates, two-way ANOVA with Tukey's test.

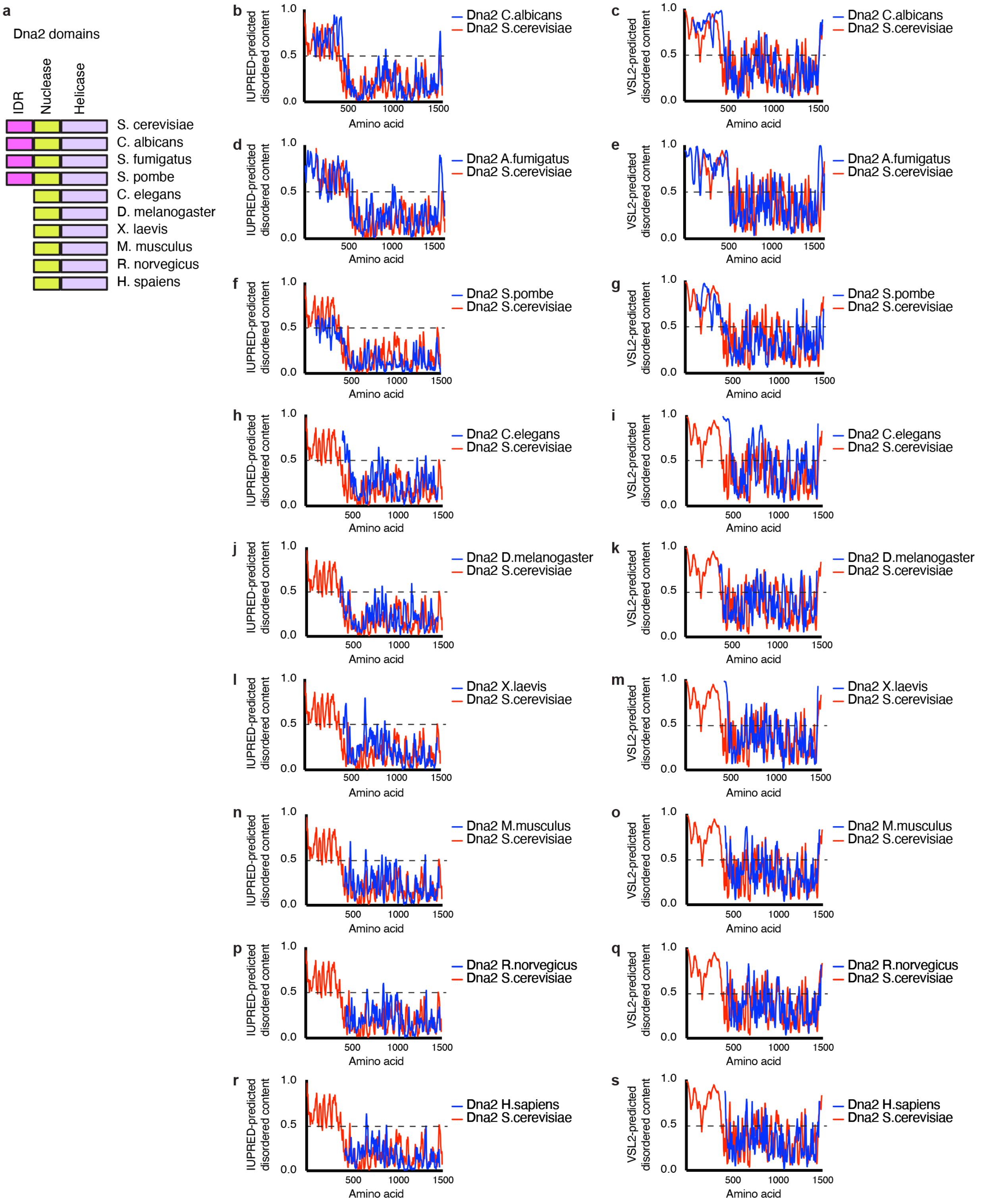


**Supplementary Fig. 8. The intrinsically disordered N-terminus of Dna2 is conserved in fungi but absent from metazoan orthologues. a**, Schematics of Dna2 protein domains in the indicated fungal and metazoan species. **b**-**s**, IUPred-predicted and PONDR-VSL2-predicted disorder profiles across the Dna2 protein in fungi (**b**-**g**) and metazoans (**h**-**s**).

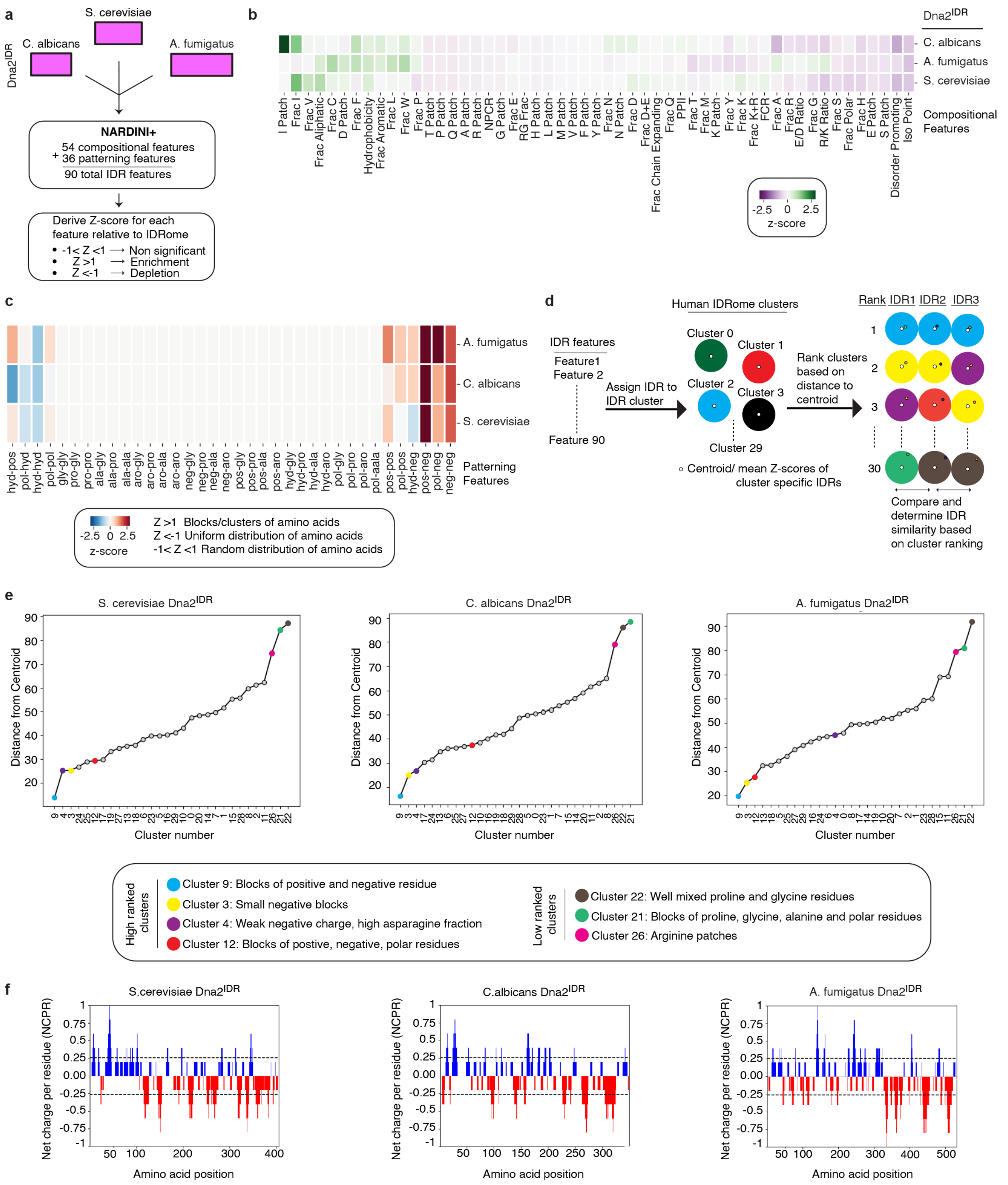


**Supplementary Fig. 9. Machine-learning-based analysis of Dna2 intrinsic disorder regions among fungi and mapping to the human IDRome. a**, Schematic of the NARDINI+ pipeline used to extract 54 compositional and 36 patterning features for the Dna2 IDR from *S. cerevisiae*, *C. albicans,* and *A. fumigatus*, and to derive feature-wise Z-scores relative to the *S. cerevisiae* IDRome reference[44]. **b,** Heat map of compositional features showing enrichment or depletion of amino-acid classes and sequence properties in each fungal

Dna2 IDR. **c**, Heat map of patterning features indicating the tendency for residues to form blocks/clusters, a uniform distribution, or a random distribution. **d**, Workflow for assigning each fungal Dna2 IDR to IDR clusters within the human IDRome by computing mean Z-scores across all 90 features, determining the nearest cluster centroids, and ranking the three best-matching IDRome clusters for each fungal IDR. **e**, Distance of each fungal Dna2 IDR to the centroids of all human IDRome clusters, with clusters coloured to highlight the closest matches that share specific sequence organizational themes. **f**, Net charge per residue (NCPR) profiles along the primary sequence of each fungal Dna2 IDR.

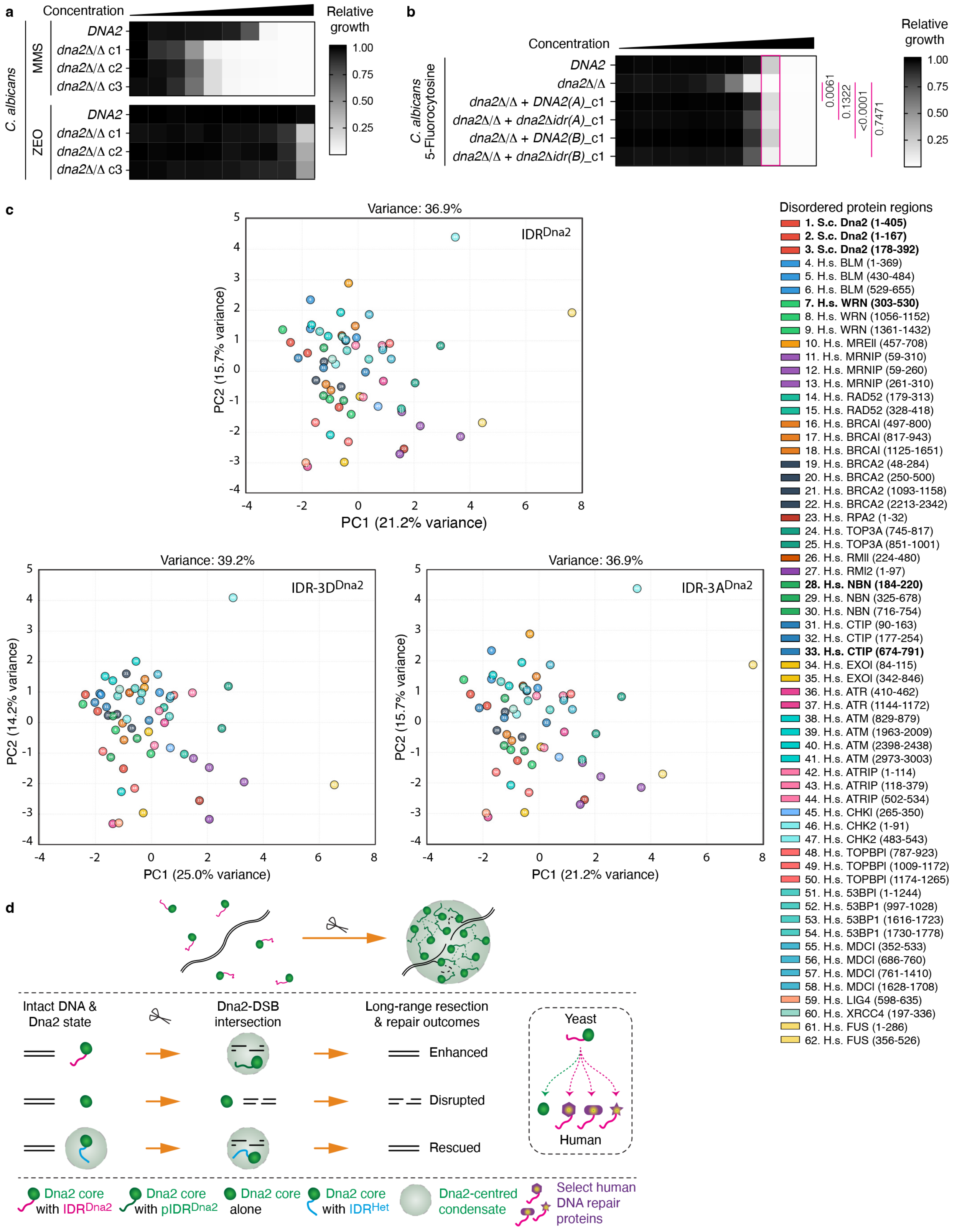


**Supplementary Fig. 10. Fungal Dna2 function and machine-learning analysis support the evolutionary redistribution of IDR$^{Dna2}$ condensate grammar. a,** Sensitivity test of wild-type and DNA2 knockout (*dna2Δ/Δ*) *C. albicans* cells to methyl methanesulfonate (MMS) or zeocin (ZEO). Both DNA2 alleles (A and B) were knocked out (*dna2Δ/Δ*), and three clones (c1-c3) were tested. The data suggest naturally higher sensitivity of *C. Albicans* to MMS compared to ZEO. **b**, Effect of removing the Dna2 IDR in *Candida albicans* Dna2 on sensitivity to the antifungal agent 5-fluorocytosine. Both *DNA2* alleles (A and B) were knocked out (*dna2Δ/Δ*) before the cells were complemented with each full-length or IDR-less Dna2. Heatmap shows the mean of *n* = 3 independent biological replicates, two-way ANOVA with Tukey's test. **c**, Principal component analysis (PCA) projections from machine-learning-based analysis assessing the similarity between the molecular grammar of *S. cerevisiae* $IDR^{Dna2}$ and intrinsically disordered regions from 25 major human DNA repair protein monomers (top plot). In this boutique analysis, $IDR^{Dna2}$ was used both in full and split into two segments according to predicted disorder, and using a similar approach, one to four highly disordered regions for each human protein analyzed were included. Numbered circles on the plot correspond to the protein and disordered amino acid ranges indicated in the legend (right; protein names and regions of particular interest are bolded). Protein monomers were analyzed without artificially filtering out conditionally structured regions to avoid bias. Also shown are versions of the same analysis, replacing $IDR^{Dna2}$ with IDR-$3D^{Dna2}$ (bottom-left plot) and IDR-$3A^{Dna2}$ (bottom-right plot). Note the closer association of the full-length IDR-$3D^{Dna2}$ variant with an additional unstructured domain of human CtIP. Also note how the $IDR^{FUS}$ region (#61, 1-286) used in our rescue experiments is different from the $IDR^{Dna2}$ (#1, 1-405). **d**, Model illustrating enzyme-intrinsic condensation in DNA end resection. Condensation intrinsic to the DNA end-processing enzyme Dna2 promotes long-range resection in yeast and is modulated by phosphorylation within its intrinsically disordered region ($IDR^{Dna2}$) (top). The effects of different Dna2 variants (namely, wild-type and $IDR^{Dna2}$-deficient Dna2, either alone or synthetically fused to the heterologous $IDR^{FUS}$) on biomolecular condensation, resection, and repair are illustrated (middle). Comparative analyses are consistent with an evolutionary redistribution of condensation-promoting features associated with $IDR^{Dna2}$ among select human DNA2-associated factors (inset). For simplicity, higher-order ssDNA organization by $IDR^{Dna2}$ condensates in the presence of yRPA and the Sgs1-dependent localization of synthetic Dna2 condensates to DNA damage sites are not illustrated. Scissors, DNA double-strand break; p, phosphorylation; Het, heterologous.

**Supplementary Table 1. *S. cerevisiae* strains used in this study.**

| Strain Nb. | Detailed genotype | Source |
|---|---|---|
| KMY372 | *(=BY4741) MATa his3Δ1 leu2Δ0 met15Δ0 ura3Δ0* | Prior study[21] |
| KMY3816 | *MATa his3Δ1 leu2Δ0 met15Δ0 ura3Δ0 DNA2-GFP-HIS3 NUP49-mCherry-HPH pDNA2-GFP-KAN* | This study |
| KMY3809 | *MATa his3Δ1 leu2Δ0 met15Δ0 ura3Δ0 pDNA2-GFP-KAN* | This study |
| KMY3891 | *MATa his3Δ1 leu2Δ0 met15Δ0 ura3Δ0 pdna2Δ405-GFP-KAN* | This study |
| KMY3892 | *MATa his3Δ1 leu2Δ0 met15Δ0 ura3Δ0 pIDRFUS-dna2Δ405-GFP-KAN* | This study |
| KMY3834 | *MATa his3Δ1 leu2Δ0 met15Δ0 ura3Δ0 pif1-m2* | This study |
| KMY3845 | *MATa his3Δ1 leu2Δ0 met15Δ0 ura3Δ0 pif1-m2 dna2Δ::HPH* | This study |
| KMY3856 | *MATa his3Δ1 leu2Δ0 met15Δ0 ura3Δ0 pif1-m2 dna2Δ::HPH pDNA2-GFP* | This study |
| KMY3857 | *MATa his3Δ1 leu2Δ0 met15Δ0 ura3Δ0 pif1-m2 dna2Δ:: HPH pdna2Δ405-GFP* | This study |
| KMY3893 | *MATa his3Δ1 leu2Δ0 met15Δ0 ura3Δ0 pif1-m2 dna2Δ::HPH pIDRFUS-dna2Δ405-GFP* | This study |
| KMY 2531 | (=TGI354) *hoΔ MATa-inc arg5,6::MATa-HPH ade3::GAL-HO hmrΔ::ADE hmlΔ::ADE1 ura3-52* | Prior study[61] |
| KMY3876 | *hoΔ MATa-inc arg5,6::MATa-HPH ade3::GAL-HO hmrΔ::ADE hmlΔ::ADE1 ura3-52 dna2(T4D,S17D,S237D)* | This study |
| KMY3874 | *hoΔ MATa-inc arg5,6::MATa-HPH ade3::GAL-HO hmrΔ::ADE hmlΔ::ADE1 ura3-52 dna2(T4A,S17A,S237A)* | This study |
| KMY3877 | *hoΔ MATa-inc arg5,6::MATa-HPH ade3::GAL-HO hmrΔ::ADE hmlΔ::ADE1 ura3-52 Dna2-GFP* | This study |
| KMY3878 | *hoΔ MATa-inc arg5,6::MATa-HPH ade3::GAL-HO hmrΔ::ADE hmlΔ::ADE1 ura3-52 dna2(T4D,S17D,S237D)-GFP* | This study |

| | | |
|---|---|---|
| KMY3879 | *hoΔ MATa-inc arg5,6::MATa-HPH ade3::GAL-HO hmrΔ::ADE hmlΔ::ADE1 ura3-52 dna2(T4A,S17A,S237A)-GFP* | This study |
| KMY3887 | *hoΔ MATa-inc arg5,6::MATa-HPH ade3::GAL-HO hmrΔ::ADE hmlΔ::ADE1 ura3-52 dna2(T4D,S17D,S237D) exo1Δ::KAN* | This study |
| KMY3884 | *hoΔ MATa-inc arg5,6::MATa-HPH ade3::GAL-HO hmrΔ::ADE hmlΔ::ADE1 ura3-52 dna2(T4A,S17A,S237A) exo1Δ::KAN* | This study |
| KMY3882 | *hoΔ MATa-inc arg5,6::MATa-HPH ade3::GAL-HO hmrΔ::ADE hmlΔ::ADE1 ura3-52 exo1Δ::KAN* | This study |
| KMY995 | *MATα his3Δ1 leu2Δ0 ura3Δ0* | This study |
| KMY3894 | *MATα his3Δ1 leu2Δ0 ura3Δ0 RFA1-CFP-KAN + pIDRFUS-dna2Δ405-GFP-KAN* | This study |
| KMY3895 | *MATα his3Δ1 leu2Δ0 ura3Δ0 RFA1-CFP-KAN + pIDRFUS-dna2Δ405-GFP-KAN sgs1Δ::HPH* | This study |
| KMY3783 | *MATa his3Δ1 leu2Δ0 met15Δ0 ura3Δ0 DNA2-GFP-HIS3* | Prior study[85] |
| KMY3542 | *MATa his3Δ1 leu2Δ0 met15Δ0 ura3Δ0 RAD52-GFP-HIS3* | Prior study[85] |
| KMY3545 | *MATa his3Δ1 leu2Δ0 met15Δ0 ura3Δ0 RFA1-GFP-HIS3* | Prior study[85] |
| KMY3792 | *MATa his3Δ1 leu2Δ0 met15Δ0 ura3Δ0 RFA2-GFP-HIS3* | Prior study[85] |
| KMY3785 | *MATa his3Δ1 leu2Δ0 met15Δ0 ura3Δ0 MRE11-GFP-HIS3* | Prior study[85] |
| KMY3780 | *MATa his3Δ1 leu2Δ0 met15Δ0 ura3Δ0 RAD59-GFP-HIS3* | Prior study[85] |
| KMY3790 | *MATa his3Δ1 leu2Δ0 met15Δ0 ura3Δ0 RTT107-GFP-HIS3* | Prior study[85] |
| KMY3775 | *MATa his3Δ1 leu2Δ0 met15Δ0 ura3Δ0 DDC2-GFP-HIS3* | Prior study[85] |

**Supplementary Table 2. *C. albicans* strains used in this study.**

| Strain Nb. | Strain details | Genotype | Parental Strain | Source |
|---|---|---|---|---|
| SC5314 | *DNA2* | *C. albicans* reference strain | - | Prior study[86] |
| CaLC11128 | *dna2Δ/Δ* c1 | *Δdna2::FRT/Δdna2::FRT* | SC5314 | This study |
| CaLC11129 | *dna2Δ/Δ* c2 | as CaLC11128 | SC5314 | This study |
| CaLC11130 | *dna2Δ/Δ* c3 | as CaLC11128 | SC5314 | This study |
| CaLC11134 | *dna2Δ/Δ+DNA2(A)* c1 | *(Δdna2::FRT)::DNA2(A)-FRT/ (Δdna2::FRT)::DNA2(A)-FRT* | CaLC11128 | This study |
| CaLC11135 | *dna2Δ/Δ+DNA2(A)* c2 | as CaLC11134 | CaLC11128 | This study |
| CaLC11136 | *dna2Δ/Δ+DNA2(A)* c3 | as CaLC11134 | CaLC11128 | This study |
| CaLC11137 | *dna2Δ/Δ+DNA2(B)* c1 | *(Δdna2::FRT)::DNA2(B)-FRT/ (Δdna2::FRT)::DNA2(B)-FRT* | CaLC11128 | This study |
| CaLC11138 | *dna2Δ/Δ+DNA2(B)* c2 | as CaLC11137 | CaLC11128 | This study |
| CaLC11139 | *dna2Δ/Δ+DNA2(B)* c3 | as CaLC11137 | CaLC11128 | This study |
| CaLC11140 | *dna2Δ/Δ+dna2Δidr(A)* c1 | *(Δdna2::FRT)::dna2Δidr(A)-FRT/ (Δdna2::FRT)::dna2Δidr(A)-FRT* | CaLC11128 | This study |
| CaLC11141 | *dna2Δ/Δ+dna2Δidr(A)* c2 | as CaLC11140 | CaLC11128 | This study |
| CaLC11142 | *dna2Δ/Δ+dna2Δidr(A)* c3 | as CaLC11140 | CaLC11128 | This study |
| CaLC11143 | *dna2Δ/Δ+dna2Δidr(B)* c1 | *(Δdna2::FRT)::dna2Δidr(B)-FRT/ (Δdna2::FRT)::dna2Δidr(B)-FRT* | CaLC11128 | This study |
| CaLC11144 | *dna2Δ/Δ+dna2Δidr(B)* c2 | as CaLC11143 | CaLC11128 | This study |
| CaLC11145 | *dna2Δ/Δ+dna2Δidr(B)* c3 | as CaLC11143 | CaLC11128 | This study |

**Supplementary Table 3. Antibodies used in this study.**

| Antibody | Company | Catalogue Nb. | Experiment |
|---|---|---|---|
| Rad53 (Rb) | Abcam | Ab104232 | WB (1:1,000) |
| Phospho-RFA2(Ser122) (Rb) | Thermo Fisher Scientific | 600-401-447 | WB (1:1,000) |
| Gamma H2A(S129) (Ms) | Abcam | Ab15083 | WB (1:1,000) |
| Beta actin (Ms) | Thermo Fisher Scientific | MA1-744 | WB (1:1,000) |
| GFP (Ms) | Thermo Fisher Scientific | A-11120 | WB (1:1,000) |
| HRP-mouse-IgG | Amersham | NA931-1ML | WB (1:10,000) |
| HRP-rabbit-IgG | Amersham | NA934-1ML | WB (1:10,000) |

**Supplementary Table 4. Primers, sgRNAs, and labelled DNA used in this study.**

| Primers | Sequence | Application |
|---|---|---|
| **For *S. cerevisiae*** | | |
| T4A, S17A HDR template FWD | TGACAATTGAAGAGATCGTCAGGATGCCCGGAG CGCCACAGAAGAACAAGAGATCTGCGAGTATAT CTGTTGCACCTGCG | Forward primer oligonucleotide for introducing T4A,S17A mutation |
| T4A, S17A HDR template RVS | TTGCTTAGATAATATCGCCTTTGAATCATTTTG TATTATTTCTTTTTCCTCTGTCTTCTTCGCAGG TGCAACAGATATAC | Reverse primer oligonucleotide for introducing T4A and S17A mutation |
| T4D, S17D HDR template FWD | TGACAATTGAAGAGATCGTCAGGATGCCCGGAG ATCCACAGAAGAACAAGAGATCTGCGAGTATAT CTGTTGATCCTGCG | Forward primer oligonucleotide for introducing T4D and S17D mutation |
| T4D, S17D HDR template RVS | TTGCTTAGATAATATCGCCTTTGAATCATTTTG TATTATTTCTTTTTCCTCTGTCTTCTTCGCAGG ATCAACAGATATAC | Reverse primer oligonucleotide for introducing T4D and S17D mutation |
| gRNA Oligo 1 T4, S17 | CTTTGAACGCCACAGAAGAACAAG | gRNA sequence specific to the region containing Threonine 4 and Serine 17 sites |
| gRNA Oligo 2 T4, S17 | AAACCTTGTTCTTCTGTGGCGTTC | gRNA sequence specific to the region containing Threonine 4 and Serine 17 sites |
| S237A HDR template FWD | GATGATATAGAGGGCGATTTAACTATAAAACCG ACGATAACGAAATTCAGCGATTTGCCATCTGCA CCCATCAAAGCACCCAACGTTGAAAAAAAAGCA GAGGTGAAT | Forward primer oligonucleotide for introducing S237A mutation |
| S237D HDR template FWD | GATGATATAGAGGGCGATTTAACTATAAAACCG ACGATAACGAAATTCAGCGATTTGCCATCTGAT CCCATCAAAGCACCCAACGTTGAAAAAAAAGCA GAGGTGAAT | Forward primer oligonucleotide for introducing S237D mutation |

| | | |
|---|---|---|
| S237A/D HDR template FWD | AGTTAAAATATCTATCAAAGAGTCATCGCCATCGTTGCTATCTCCTGTTGAATCCATTTTATCTACTTCTTCTGCATTCACCTCTGCTTTTTTTTCAACGTTGGG | Reverse primer oligonucleotide for introducing either S237A or S237D mutation |
| gRNA Oligo 1 S237 | AAACTGATGGGTGAAGATGGCAAA | gRNA sequence specific to the region containing the Serine 237 site |
| gRNA Oligo 2 S237 | CTTTTTTGCCATCTTCACCCATCA | gRNA sequence specific to the region containing the Serine 237 site |
| 3.4 Kb Downstream of HO cut FWD | GTACTCCCGTCGTGTCTTATC | qPCR primer to measure resection |
| 3.4 Kb downstream of HO cut RVS | ATACCTAAATCAACCAAGTCACG | qPCR primer to measure resection |
| 6.6 Kb Downstream of HO cut FWD | AAGCCAAAGAAGAAGAAATGCT | qPCR primer to measure resection |
| 6.6 Kb Downstream of HO cut RVS | AGACGTAACAAGCTTCTAGTGAATA | qPCR primer to measure resection |
| ACT1 FWD | GCCTTCTACGTTTCCATCCA | qPCR primer control for resection measurement (from prior study[62]) |
| ACT1 RVS | GGCCAAATCGATTCTCAAAA | qPCR primer control for resection measurement (from prior study[62]) |
| **For recombinant *S. cerevisiae* Dna2** | | |
| oKM1001 | 5′-Cy5-ACGCTGCCGAATTCTACCAGTGCCTTGCTAGGACATCTTTGCCCACCTGCAGGTTCACCC-3′ | Oligo 1 of 3 for creating the 5’-flap substrate for *S.c.* Dna2 in vitro nuclease assays and imaging |

| | | |
|---|---|---|
| oKM1002 | 5′-GGGTGAACCTGCAGGTGGGCAAAGATGTCCATCTGTTGTAATCGTCAAGCTTTATGCCGT-3′ | Oligo 2 of 3 for creating the 5'-flap substrate for *S.c.* Dna2 in vitro nuclease assays and imaging |
| oKM1003 | 5′-GGGTGAACCTGCAGGTGGGCAAAGATGTCCCATGGAGCTGTCTAGAGGATCCGACTATCG-3′ | Oligo 3 of 3 for creating the 5'-flap substrate for *S. c.* Dna2 in vitro nuclease assays and imaging |
| oKM1004 | 5′-Cy3-ATGTGGAACAGTGGATTCGAAAGCTATGGCAGCGACGACT-3′ | Oligo representing the ssDNA molecule used for imaging-based experiments |
| **For *C. albicans*** | | |
| oLC14458 | CCGTGAACCAATGCGTTGACCAAATTAAAAATAGTTTACGCAAGTC | For generating sgRNA to delete *CaDNA2* |
| oLC14459 | GTCAACGCATTGGTTCACGGGTTTTAGAGCTAGAAATAGCAAGTTAAA | For generating sgRNA to delete *CaDNA2* |
| oLC6926 | AAGAAAGAAAGAAAACCAGGAGTGAA | Universal primer for producing sgRNA (from prior study[78]) |
| oLC6927 | ACAAATATTTAAACTCGGGACCTGG | Universal primer for producing sgRNA (from prior study[78]) |
| oLC6928 | GCGGCCGCAAGTGATTAGACT | Universal primer for producing sgRNA (from prior study[78]) |
| oLC6929 | GCAGCTCAGTGATTAAGAGTAAAGATGG | Universal primer for producing sgRNA (from prior study[78]) |
| oLC14460 | ATGTGATATCTCATTGATAGTCGAAGAGGTAGTAGCTACACACTTGCTTTTGTGCTTGGAATCAAACGCGCTGAAATCGGAGGAAACAGCTATGACCATG | Primer with upstream homologous region to amplify the *NAT-FLP* cassette for deleting *CaDNA2* |
| oLC14461 | AGCTATAGCTAACTACAAACAATGCATAACCAATCTCTAAAAATTTTCTAGTTTATCTAGTAATGT | Primer with downstream homologous region to amplify the *NAT-FLP* cassette for deleting *CaDNA2* |

| | | |
|---|---|---|
| | CTTGAACCACATTCGTAAAACGACGGCCAGTGAG | |
| oLC14462 | GCAACTGGAACTAAACTTTCTCCAGGAG | Forward primer for checking the absence of *CaDNA2* coding sequence |
| oLC14463 | GGTAACAGGTTTGGATTTTGGTCG | Reverse primer for checking the absence of *CaDNA2* coding sequence |
| oLC11150 | CCGCGGTGGAGCTCCAATTCCAAATTAAAAATAGTTTACGCAAGTC | Primer to generate sgRNA for re-introducing *CaDNA2* alleles to the deletion mutant |
| oLC11151 | GAATTGGAGCTCCACCGCGGGTTTTAGAGCTAGAAATAGCAAGTTAAA | Primer to generate sgRNA for re-introducing *CaDNA2* alleles to the deletion mutant |
| oLC14469 | GACAAATGGGACAGAGCTCCATGGC | Forward primer for confirming re-introduction at both *CaDNA2* loci |
| oLC14470 | GAAGAATGAGAAACGGTCAACGGAGAG | Forward primer for confirming re-introduction at both *CaDNA2* loci |
| oLC14464 | CATTACTCTCGAGACTAGAAAATTTTTAGAGATTGGTTATGCATTGTTTGTAGTTAGCTATAGCTCTATATAGCAAAGAAATGAAATTCAAAATTACCTC | Forward primer with an XhoI site for amplifying the *CaDNA2* 3’ downstream region |
| oLC14465 | TGACGGGCCCGATGACAATCAAGGGCAAAATGGCTGGAATGGTAATCAATTCCATTTCAAAAGAGGTAATTTTGAATTTCATTTCTTTGCTATATAGAGC | Reverse primer with an ApaI site for amplifying the *CaDNA2* 3’ downstream region |
| oLC14466 | GATTGGAGCTCCGAGTCTTTATACGGTTGACTATATTTGG | Forward primer with a SacI site for amplifying the *CaDNA2* 5’ upstream and coding region |
| oLC14467 | ATCTCCCGCGGTTTTCTAGTTTATCTAGTAATGTCTTGAACCACATTCTTAAGTATAGGG | Reverse primer with a SacII site for amplifying the *CaDNA2* 5’ upstream and coding region |

| | | |
|---|---|---|
| oLC14471 | CTACTTTTATATCAAACACTTATCATAGTatgG GTATTCAAAACTTTGAAAAACTAAATATTG | Fusion PCR primer for truncating *CaDNA2 IDR* |
| oLC14472 | ACTATGATAAGTGTTTGATATAAAAGTAG | Fusion PCR primer for truncating *CaDNA2 IDR* |

**Supplementary Table 5. Plasmids used in this study.**

| Plasmid Nb. | Details | Source |
|---|---|---|
| **For *S. cerevisiae*** | | |
| pKM718 | *pdna2Δ405-GFP-KANMX6* | This study |
| pKM731 | *pIDRFUS-dna2Δ405-GFP-KANMX6* | This study |
| pKM541 | *pDNA2-GFP-KANMX6* | Previous study[48] |
| pKM190 | *pGFP-TUB1-URA3* | Previous study[87] |
| pKM734 | *pCAS* | Addgene (Cat. Nb. 60847)[88] |
| **For *C. albicans*** | | |
| pV1093 | Tool vector for transient CRISPR | Prior studies[78,89] |
| *pSFS2-SAT1* | *SAT1* flipper | Prior study[67] |
| pLC49 | *NAT* flipper | Prior study[79] |
| pLC2112 | *pSFS2-DNA2(A)* | This study |
| pLC2113 | *pSFS2-DNA2(B)* | This study |
| pLC2115 | *pSFS2-dna2Δidr(A)* | This study |
| pLC2116 | *pSFS2-dna2Δidr(B)* | This study |

### Supplementary Video Legends

**Supplementary Video 1. Dna2 repair foci exhibit kinetics consistent with liquid-like behavior.** Dna2-GFP repair foci exhibit rapid signal recovery in fluorescence recovery after photobleaching (FRAP). DNA damage was induced using methyl methane sulfonate (MMS; 0.03%). A magenta-colored circle indicates the photobleached focus. Scale bar, 0.5 µm.

**Supplementary Video 2. Liquid-like fusion of IDR$^{Dna2}$ droplets in vitro.** GFP-IDR$^{Dna2}$ droplets undergo efficient fusion followed by relaxation into a relatively more spherical shape. Shown are the GFP and DIC views of the fusing droplets. Scale bar, 1 µm.

**Supplementary Video 3. IDR$^{Dna2}$ droplets show internal kinetics consistent with liquid-like behavior.** GFP-IDR$^{Dna2}$ droplets show rapid signal recovery in fluorescence recovery after photobleaching (FRAP). Scale bar, 2 µm.

**Supplementary Video 4. Reorganization of ssDNA by mature IDR$^{Dna2}$ assemblies in the presence of yRPA.** A magenta oval is used to temporarily highlight Cy3-ssDNA-positive filament dynamics and its crosstalk with GFP-IDR$^{Dna2}$ assemblies. Imaging was conducted after a 15 min incubation of GFP-IDR$^{Dna2}$ with yRPA, followed by a 60 min incubation with Cy3-ssDNA. Scale bar, 2 µm.
